# Structural analysis and conformational dynamics of a holo-adhesion GPCR reveal interplay between extracellular and transmembrane domains

**DOI:** 10.1101/2024.02.25.581807

**Authors:** Szymon P. Kordon, Kristina Cechova, Sumit J. Bandekar, Katherine Leon, Przemysław Dutka, Gracie Siffer, Anthony A. Kossiakoff, Reza Vafabakhsh, Demet Araç

## Abstract

Adhesion G Protein-Coupled Receptors (aGPCRs) are key cell-adhesion molecules involved in numerous physiological functions. aGPCRs have large multi-domain extracellular regions (ECR) containing a conserved GAIN domain that precedes their seven-pass transmembrane domain (7TM). Ligand binding and mechanical force applied on the ECR regulate receptor function. However, how the ECR communicates with the 7TM remains elusive, because the relative orientation and dynamics of the ECR and 7TM within a holoreceptor is unclear. Here, we describe the cryo-EM reconstruction of an aGPCR, Latrophilin3/ADGRL3, and reveal that the GAIN domain adopts a parallel orientation to the membrane and has constrained movement. Single-molecule FRET experiments unveil three slow-exchanging FRET states of the ECR relative to the 7TM within the holoreceptor. GAIN-targeted antibodies, and cancer-associated mutations at the GAIN-7TM interface, alter FRET states, cryo-EM conformations, and receptor signaling. Altogether, this data demonstrates conformational and functional coupling between the ECR and 7TM, suggesting an ECR-mediated mechanism of aGPCR activation.

## INTRODUCTION

With 33 members in humans, the adhesion G protein-coupled receptors (aGPCRs) make up the second largest GPCR family, but the molecular mechanisms underlying their activation and modulation are not fully understood^1–3^. Genetic studies have demonstrated critical roles for aGPCRs in development, immunity, and neurobiology, including brain development^4–8^, myelination^9^, brain angiogenesis^10^ and neural tube development^11,12^. They are also linked to various diseases such as neurodevelopmental disorders, deafness, male infertility, attention deficit hyperactivity disorder, schizophrenia, immune disorders, and cancers^6,13–16^. While 30% of FDA-approved drugs target GPCRs^17^, aGPCRs have yet to be targeted therapeutically, primarily due to our limited understanding of their functional modulation.

The distinctive chimeric architecture of aGPCRs differentiates them from conventional GPCRs^13,18^. In addition to their signaling seven transmembrane (7TM) helices that are characteristic of all GPCRs, aGPCRs also have large (up to 6000 residues), multidomain extracellular regions (ECRs). The ECR is characterized by a conserved GPCR Autoproteolysis INducing (GAIN) domain that is always positioned at the far C-terminus of the ECR, in close proximity to the 7TM region^19^. The GAIN domain contains an autoproteolysis site between its last two β-strands and it is cleaved during protein maturation in the endoplasmic reticulum^19–21^. After autoproteolysis, the cleaved fragments stay non-covalently connected on the cell surface. The multidomain structure of the ECR allows it to interact with protein ligands found on adjacent cells or within the extracellular matrix. These interactions give rise to mechanical forces that are key for regulating receptor activation by modulating the accessibility of the last beta strand of the GAIN domain, that is also called the tethered agonist (TA) or Stachel peptide. Under normal conditions, the TA peptide lies within the core of the GAIN domain^22,23^, but upon force-induced dissociation of the cleavage fragments, the TA peptide becomes exposed and can bind to the orthosteric site of the 7TM domain to activate the receptor^24–26^. Hence, according to the TA-mediated model of aGPCR activation, the ECR acts as a protective cap for the TA peptide to hide it within the GAIN domain.

Several recent observations suggest that other mechanisms of aGPCR activation are possible^27,28^. For example, some aGPCRs do not undergo autoproteolysis, which is required for TA release^29^. Even the aGPCRs that are cleaved do not always require cleavage for mediating some aspects of wild type functions^7,29–31^. Specifically, functional data suggest that some aGPCRs are modulated by direct communication of the ECR with the 7TM^7,28,32–34^. Alterations to the ECR of aGPCRs, including alternative splicing, mutagenesis or binding of synthetic proteins have been shown to modify receptor signaling^32,35,36^. Finally, the removal of the ECR in GPR56 increased the basal activity of the receptor^37^. Furthermore, TA mechanism implies that the receptor is used only once, which would be energetically costly for the cell, and does not provide a mechanism for how the signal can be turned off when the activating ligand dissociates from the ECR. Additionally, compression forces that “push” on the receptor, for example when cells are approaching each other during synapse formation, organogenesis or embryogenesis, might not be able to use the TA-mediated mechanism to activate the aGPCR. It has been suggested that the TA can regulate receptor signaling without coming out of the GAIN domain or by being partially exposed^38^, however the recent TA-bound 7TM structures of multiple aGPCRs showed that the critical phenylalanine residue and other important TA residues reach deep into the 7TM orthosteric pocket for receptor activation, suggesting that non-release or partial release of the TA is unlikely to activate the receptor^39–42^. Thus, accumulating data in the aGPCR field are consistent with an additional model in which the ECR conformation has a direct role in modulating the 7TM signaling, independently of TA-mediated activation^27,28,37,43^.

In this ECR-mediated model for aGPCR activation, the ECR directly communicates with the 7TM (i.e. via transient interactions), such that ligand binding events or conformational changes in the ECR may directly result in altered signaling^27,28^. Direct proof of this model requires studying a holoreceptor, which is technically challenging. Several studies have determined the high-resolution structures of the GAIN domain, other extracellular domains and the 7TM from multiple aGPCRs^19,28,32,39–42,44–46^. However, none of these studies addressed how the extracellular region transduces the incoming adhesion signal to the membrane anchored 7TM domain as none of them could visualize the ECR and the 7TM simultaneously. A full-length aGPCR structure is still missing and characterization of the conformational dynamics of a full-length aGPCR remain absent.

ADGRL3, also known as Latrophilin3, is a prototypic aGPCR composed of lectin, olfactomedin, hormone binding, GAIN and 7TM domains (Fig. 1a)^47^. Previous work has shown that ADGRL3 couples to Gα_s_, Gα_12/13_, Gα_i_ and Gα ^7,48–50^. ADGRL3 is highly expressed in the brain where it is required for the establishment of excitatory synapses and for determining their specificity; and has been implicated in attention deficit-hyperactivity disorder and various cancers^7,15,51,52^. ADGRL3 mediates its synapse-formation function by its extracellular adhesive interactions with ligands, and, also, by its intracellular GPCR signaling^7,50,53–55^.

**Fig. 1:**
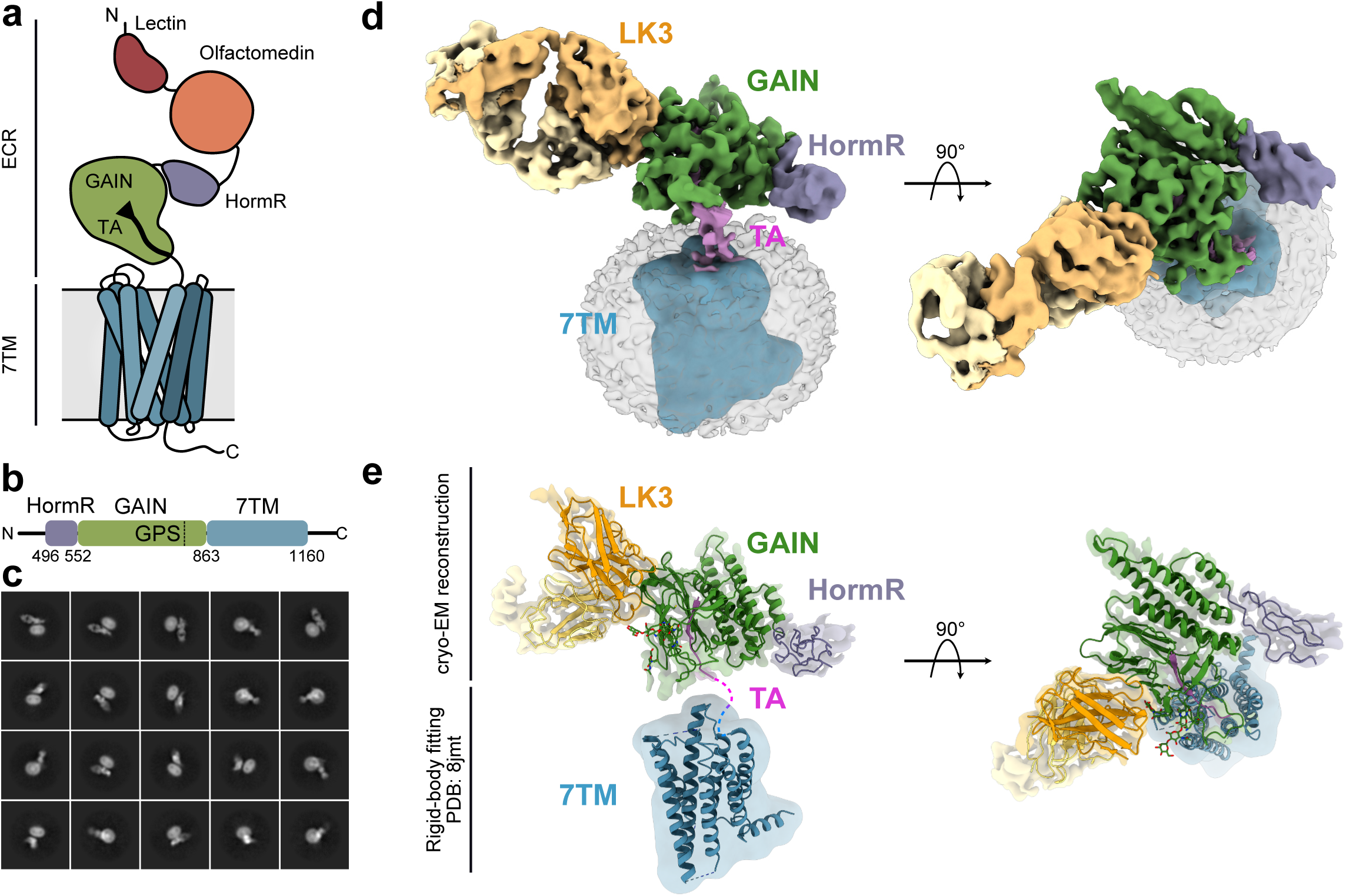
Cryo-EM model of ADGRL3/LK3 complex reveals the general architecture of the receptor and relative position of the GAIN domain in relation to 7TM and membrane. **a** Cartoon representation of ADGRL3. The lectin domain is colored red, olfactomedin - orange, HormR - purple, GAIN - green and 7TM in blue. The tethered agonist (TA) is presented as a black arrow within the GAIN domain. **b** Schematic diagram of domain boundaries of the ADGRL3 construct used for structural analysis. Domains colored as in (a) GPS represents GPCR Proteolysis Site. **c** Representative cryo-EM reference-free 2D class averages of the ADGRL3/LK3 complex purified in detergent micelle. **d** Low resolution 3D map of ADGRL3 in complex with sAB LK3. Regions of the map corresponding to respective domains of the receptor and sAB are colored accordingly and labeled with domain names. The tethered agonist (TA) is colored magenta. Density for 7TM region was Gaussian-filtered to 10 Å. **e** The composite map of ADGRL3 architecture consisting of 3.9 Å structure of the HormR/GAIN domain in complex with sAB LK3, and low-resolution density of the 7TM as in (d). Docked in the 7TM region by rigid-body fitting is the available structure of apo-state ADGRL3 corresponding to PDB 8jmt.

Here, we used a combination of single-particle cryogenic electron microscopy (cryo-EM), single-molecule Förster Resonance Energy Transfer (smFRET), protein engineering and GPCR signaling studies to map the relative orientation and conformational dynamics between the GAIN and 7TM domains of a ADGRL3 holoreceptor. Our results showed that point mutations at the GAIN-7TM interface or ECR-targeted antibodies that alter the downstream receptor signaling, also change conformational distribution of ECR in ADGRL3, demonstrating direct link between receptor activity and the orientation of the GAIN domain with respect to the 7TM domain.

## RESULTS

### Purification of ECR-bound ADGRL3 and development of synthetic antibody binders targeting the GAIN domain of ADGRL3

For structure determination we recombinantly expressed truncated human ADGRL3 containing HormR/GAIN and 7TM domains (Fig. 1b) in *Spodoptera frugiperda* Sf9 cells, solubilized from membranes using n-Decyl-beta-Maltoside (DM), and purified by affinity chromatography using FLAG-tag. DM-solubilized ADGRL3 was then subjected to size exclusion chromatography (SEC) with simultaneous detergent exchange to lauryl maltose neopentyl glycol (LMNG)/glyco-diosgenin (GDN) (Supplementary Fig. 1a). We previously reported that this construct is active in a G-protein coupling assay^24^. The sample quality was further confirmed by negative stain electron microscopy (nsEM), that showed the uniform distribution of homogenous particles of the expected shape and size (Supplementary Fig. 1b-d).

To aid the cryo-EM structural analysis of ADGRL3, we screened a synthetic antibody binder (sAB) library to obtain binders specific to the HormR/GAIN domains of ADGRL3. A biotinylated fragment of the ADGLR3 ECR containing HormR and GAIN domains was used as input for generation of high-affinity sABs against ADGRL3 by phage display selection using a diverse synthetic phage library based on a humanized antibody Fab scaffold^56–58^, and four rounds of selection were performed as described previously^35^. Following the selection and initial validation, we identified 10 unique HormR/GAIN binders by a single-point phage enzyme-linked immunosorbent assay (ELISA) (Supplementary Fig. 1e). Selected phagemids were then cloned into sAB protein format, expressed, and purified for further characterization. Two of the best expressing sABs, named LK1 and LK3 that bind to the ADGRL3 with low nanomolar affinity (3.3 and 3.9 nM respectively, Supplementary Fig. 1f), and form stable complexes with the receptor on SEC (Supplementary Fig. 1g, h) were chosen for structural and functional studies of ADGRL3.

### Cryo-EM model of ECR-bound ADGRL3 reveals the orientation of the GAIN domain with respect to the 7TM region

For structure determination of ADGRL3, we purified the ECR-bound ADGRL3 in complex with the sAB LK3 in LMNG/GDN detergent mix (Supplementary Fig. 1g). Addition of LK3 as a fiducial marker allowed for improved ADGRL3 2D classification and alignment, and enabled to obtain the overall cryo-EM projections of the complex in the micelle (Fig. 1c, Supplementary Fig. 2a). The conformational heterogeneity caused by the flexibility between the ECR and the 7TM domain restricted our attempts to obtain a high-resolution structure of the entire ADGRL3 molecule. In contrast to previous studies that utilized addition of G-proteins to lock the 7TM region in an active state^39,49^, we could not resolve the transmembrane region to high resolution. However, we can unambiguously define the position of the GAIN domain in relation to the 7TM (Fig. 1d, e). The nominal resolution of the entire complex model was limited to 5.5 Å, with most of the ECR resolved to 4 Å, and the 7TM/micelle region limited to 8-10 Å (Supplementary Fig. 2a-c, Supplementary Fig. 3a). The overall architecture of the receptor suggests that the GAIN domain of ADGRL3 lays flat, with a parallel orientation in relation to the membrane, and does not adapt the extended, vertical position as suggested by AlphaFold2 structure predictions^59^ (Supplementary Fig. 4). Though GAIN stays in a close proximity to the membrane, at our current reconstruction quality we are unable to describe any potential interaction between GAIN and 7TM regions.

To better understand the extent of the ECR/7TM flexibility in the ADGRL3 model, we performed a 3D Variability analysis and found that the ECR can sample several other conformations (Fig 2a-c)^60^. Despite this, the scope of the ECR movement retains the overall orientation of the GAIN domain, with the same side of the GAIN always facing the membrane. Comparing all conformations to the consensus model, the GAIN domain can either adopt a more upright conformation, staying right above the 7TM (Fig. 2a), or lay closer to the membrane surface, shifting laterally away from the 7TM region (Fig. 2c). However, the movement of the GAIN domain is constrained to a small volume, and does not sample the full extent of the available (calculated approximately to 30% of the total possible volume it may occupy). The magnitude of the GAIN movement between the two most distant conformations is around 45⁰ and varies between 15 Å distance change close to the 7TM/membrane and 75 Å change at the tip of the GAIN domain (Fig. 2d).

**Fig. 2:**
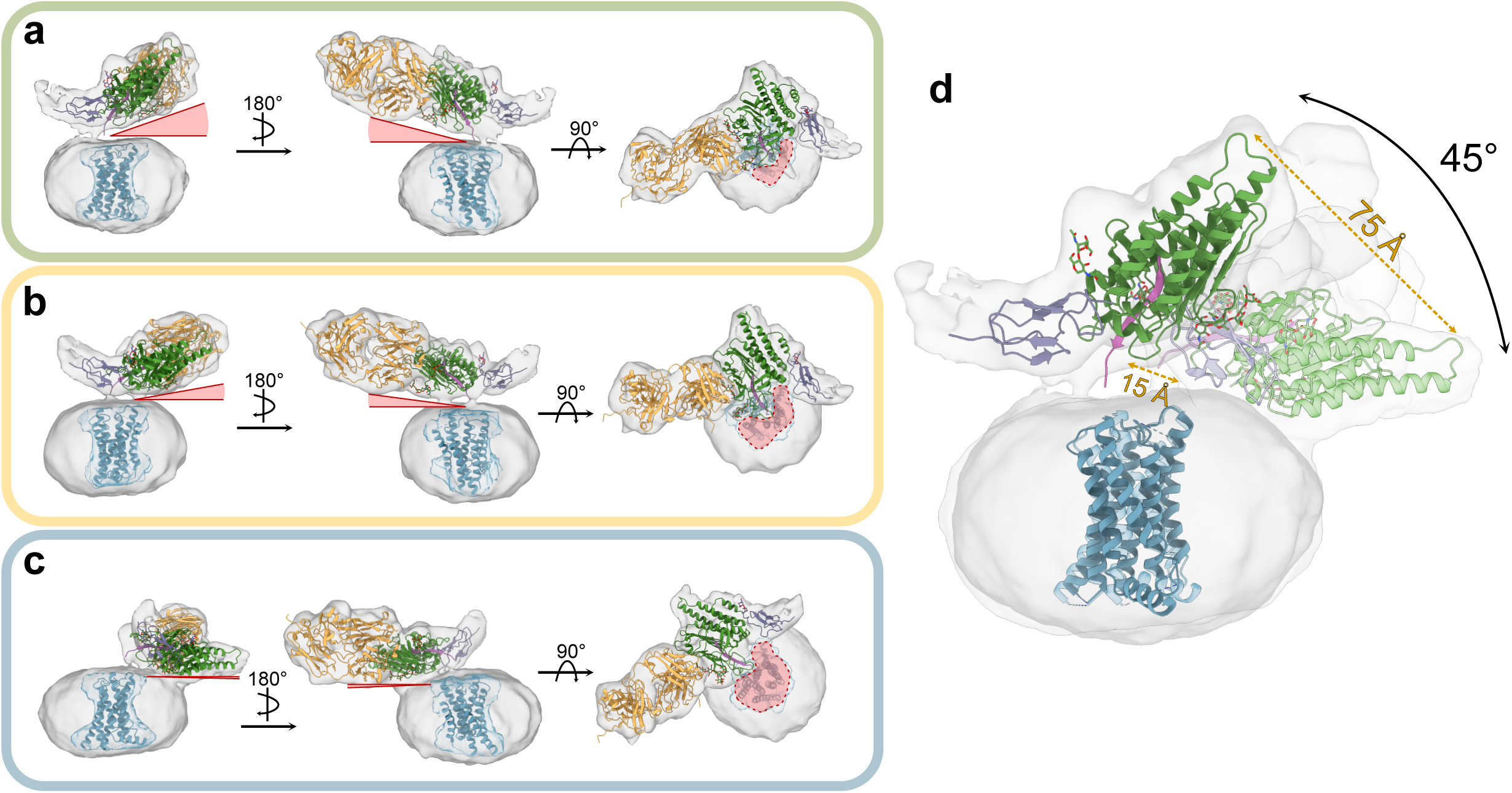
CryoEM 3D variability analysis presents constrained flexibility of the ADGRL3 GAIN domain. **a-c** Side and top views of representative low-resolution 3D models from 3D variability analysis show the degree of HormR/ GAIN domain movement in relation to the detergent micelle. Models from (a) the first frame of 3D variability analysis, showing the most vertical orientation of the GAIN domain seen within the dataset; (b) The middle frame of 3D variability analysis, presenting the intermediate position of the GAIN domain; (c) The last frame of 3D variability analysis, showing the most horizontal orientation of the GAIN domain. The red triangles on each panel depict the degree of the angle between the bottom of the GAIN domain and the plane of the micelle. The red areas highlighted on the top-view projections show the increasing access of the 7TM region to the extracellular space with the GAIN domain pulling away from the central axis. **d** Superimposition of the most distinct frames (a) and (b) showing the distances and angle between two GAIN conformations. Structures of HormR/GAIN domains of ADGRL3 in complex with LK3 and 7TM region (PDB: 8jmt) are automatically docked in the maps.

### 3.9 Å structure of HormR/GAIN domains of ADGRL3 in complex with sAB LK3

Utilizing local refinement, we determined a 3.9 Å structure of ADGRL3 HormR/GAIN domains bound to LK3^61^ (Fig. 1e, Fig. 3a). The structure shows the conserved HormR/GAIN fold, with the N-terminal subdomain A of the GAIN domain composed of six α-helices and the C-terminal subdomain B including 13 β-strands, previously reported in crystal structures of separated HormR/GAIN domains (Supplementary Fig. 5)^19,45^. Importantly, we observed a distinct density that corresponds to the TA peptide, suggesting that in holo-ADGRL3 the TA, with crucial F857, L860 and M861 in particular, stays buried in the GAIN, inaccessible to the solvent and away from the orthosteric pocket within 7TM domain, despite the considerable flexibility of the ECR (Fig. 3b). All well-resolved residues of TA peptide (T855-E865) form a strong network of hydrogen bonding, and numerous hydrophobic interactions, with total interface area of 1100 Å^2^, similar to that of isolated GAIN domains. Additionally, there is no evidence of the TA peptide coming out of, or getting partially exposed from the GAIN domain or interacting with the 7TM domain.

**Fig. 3:**
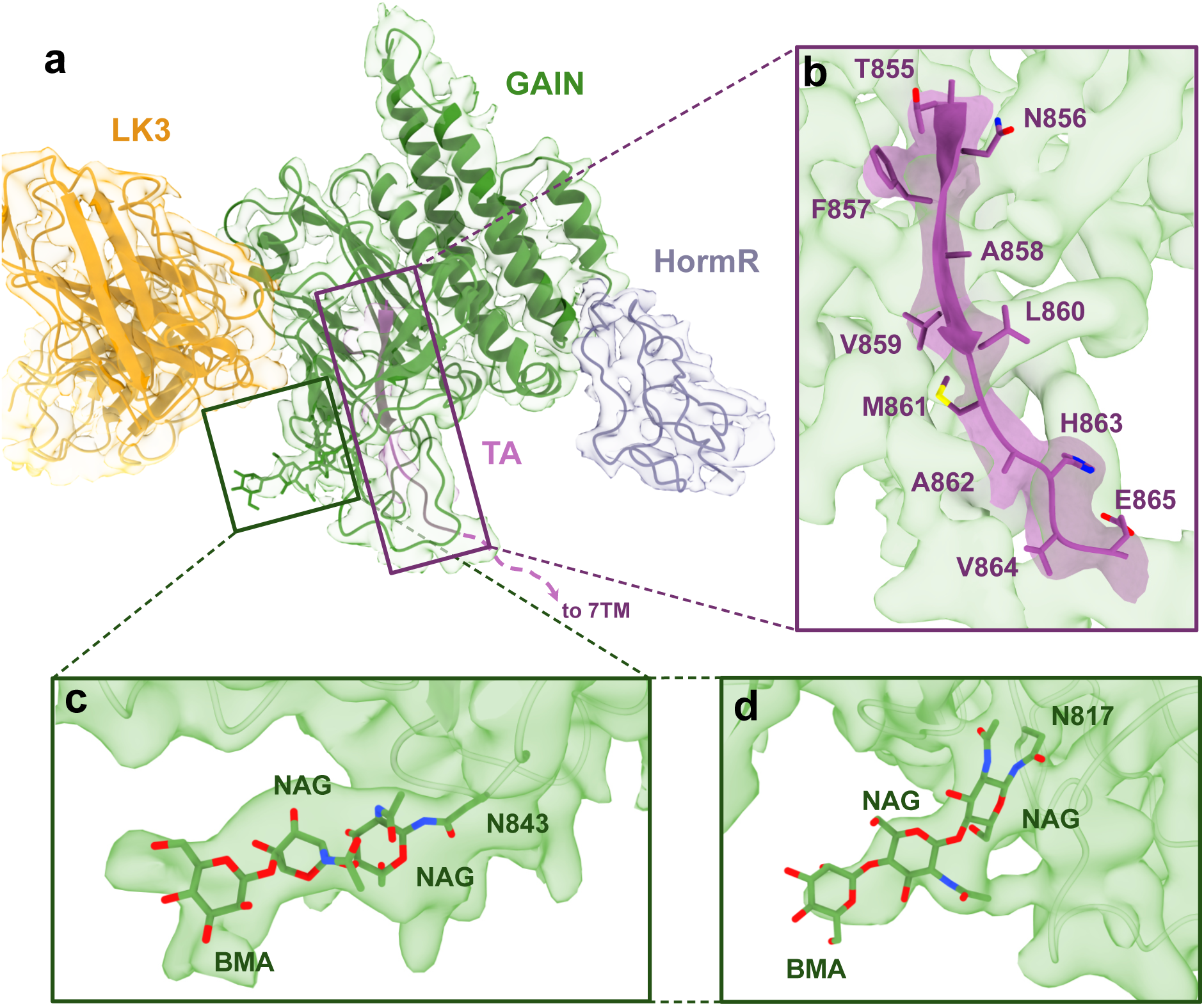
Cryo-EM structure of the 7TM-bound ADGRL3 HormR/GAIN domains at 3.9 Å resolution. **a** The cryo-EM structure and density map of HormR/GAIN domains of ADGRL3 bound to sAB LK3. HormR domain is colored purple, GAIN - green, TA - magenta, and heavy and light chains of LK3 - yellow. **b** Close-up view of the intact TA buried within the GAIN domain in the holoreceptor. Side chains of the TA peptide are shown as sticks. **c** and **d** Glycosylation of conserved sites N843 (c) and N814 (d) at the membrane-facing side of the GAIN domain modeled into the density map of HormR/GAIN domains.

Furthermore, we observed densities for at N-linked glycosylation sites on residues N843 and N817 that are conserved within ADGRLs (Fig. 3c, d). Specifically, the low-threshold map of the GAIN domain showed a density for a long glycan moiety attached at position 843, running alongside the bottom face of the GAIN domain, parallel to the membrane. It is possible that this long sugar addition may be important for maintaining the GAIN domain conformation and the orientation of the entire ECR with respect to the 7TM and the membrane, as in the case of EGFR^62^ and thyrotropin receptor ectodomains^63^. It has been shown that electrostatic interactions accompanied by steric effects between glycan chains in the protein and the membrane surface can be critical for the arrangement of extracellular domains^64^.

The cryo-EM structure also allowed us to elucidate the molecular basis of the interaction between LK3 and ADGRL3 ECR. LK3 binds to the side of the GAIN domain through CDRs in both heavy (H1, H2, H3) and light chains (L3), resulting in the total interface area of 835 Å^2^ in the complex (Supplementary Fig. 6). This interface is mediated mainly by bulky, aromatic residues of LK3 CDRs and involves extensive hydrogen bonding and van der Waals interactions with residues in subdomain B of the GAIN domain. Notably, hydrogen bonds between Y36_H1_ and A797_ADGRL3_/D798_ADGRL3_, S58_H2_ and L796_ADGRL3_/A797_ADGRL3_, Y104_H3_ and L796_ADGRL3_, W105_H3_ and S850_ADGRL3_, W108_H3_ and P799_ADGRL3_, Y112_H3_ and D798_ADGRL3_/N730_ADGRL3_, Y95_L3_ and N730_ADGRL3_, stabilize the interaction and shape the total buried surface area (Supplementary Fig. 6).

### Real-time measurement of conformational changes via smFRET present three stable conformations for GAIN/7TM

To map the conformational landscape of the full-length ADGRL3 and directly visualize the relative orientation and dynamics of the GAIN domain with respect to the 7TM domain, we used single-molecule Fӧrster resonance energy transfer (smFRET). To minimize perturbations to the receptor structure we used unnatural amino acid incorporation and click chemistry for site-specific labeling of the receptors^65,66^. With this approach we generated multiple FRET sensors to probe the relative orientation of the GAIN and the 7TM domains. Specifically, we focused on three FRET sensors with fluorophores at positions Y795UAA:R933UAA, Y795UAA:E1099UAA, and Y795UAA:A871UAA (Fig. 4a). Receptors harboring the unnatural amino acids were expressed in HEK293T cells and site-specifically labeled with FRET donor (Cy3) and acceptor (Cy5) fluorophores and showed strong labeling on the plasma membrane (Fig. 4b). After solubilization with the detergent, receptors were applied onto a polyethylene glycol (PEG)-passivated coverslips with an anti-GFP-tag antibody for total internal reflection fluorescence (TIRF) imaging^67^ (Fig 4c).

**Fig. 4:**
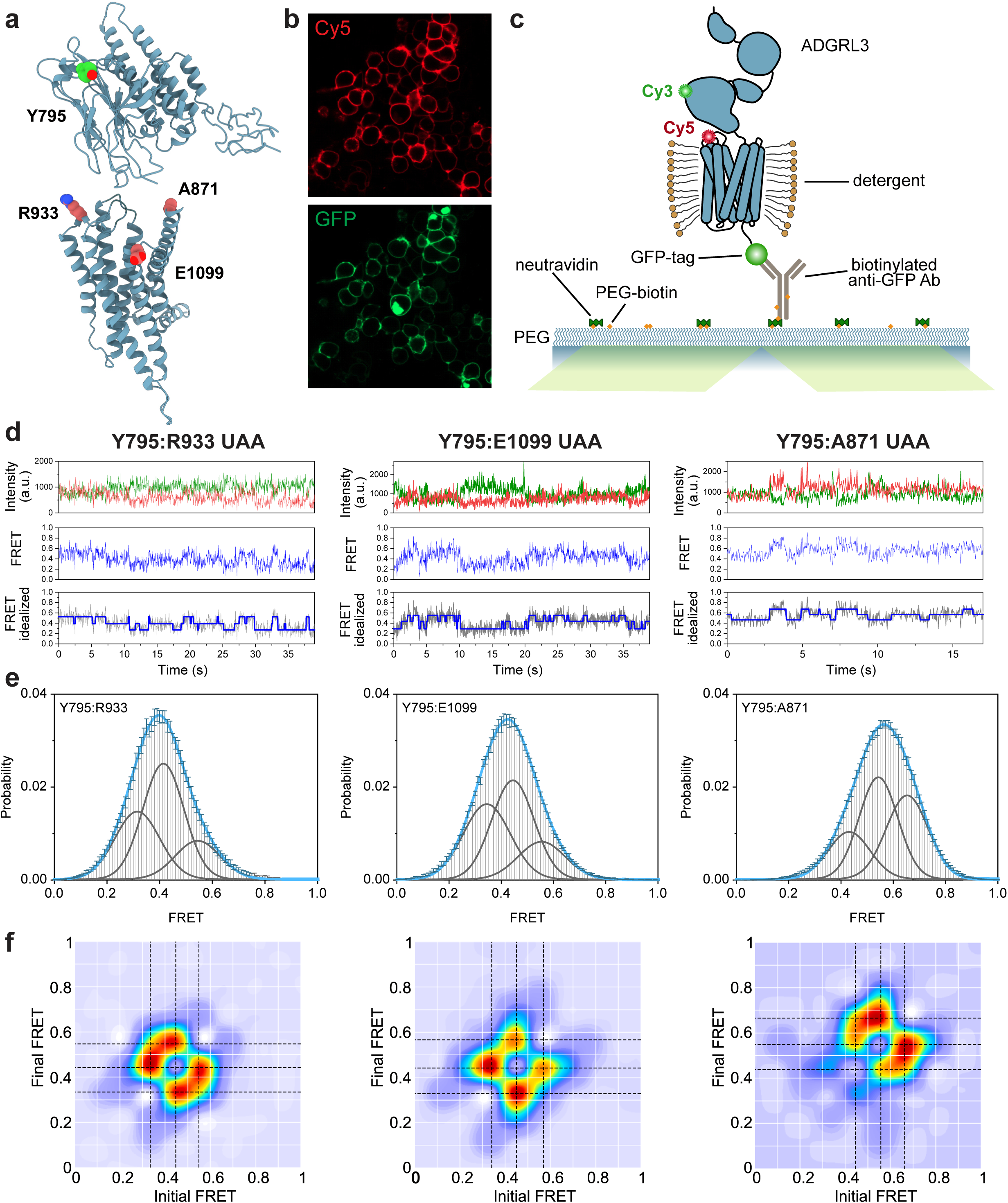
smFRET present three stable conformations of ADGRL3 ECR/7TM. **a** Model of ADGRL3 (truncated after HormR domain) showing positions of amino acids that were replaced by UAA in making of the smFRET sensor pairs: Y795 (in green), R933, E1099, A871 (in red) **b** Example of confocal microscopy images of ADGRL3-UAA labeled with Cy5 (red) showing membrane expression and GFP (green) showing overall expression of the receptor **c** Schematic of SiMPull smFRET TIRF microscopy experiment. **d** Representative smFRET traces of Y795:R933 UAA, Y795:E1099 UAA, and Y795:A871 UAA, showing donor (green) and acceptor (red) intensities, corresponding FRET (blue), and idealized FRET trace (blue line over gray FRET data) from HMM. **e** Single-molecule FRET population histograms of Y795:R933 UAA, Y795:E1099 UAA, Y795:A871 UAA, respectively (error bars represent s.e.m., *n* = 3, independent biological replicates). Three single Gaussian distributions (gray) were fitted to the histograms. Peaks are centered at 0.32, 0.41, and 0.54 for Y795:R933 UAA, 0.35, 0.44, and 0.55 for Y795:E1099 UAA, and 0.45, 0.55, 0.65 for Y795:A871 UAA. The cumulative fit is overlaid in blue. **f** Transition density plots of Y795:R933 UAA, Y795:E1099 UAA, and Y795:A871 UAA, respectively. Dashed black lines cross the most observed transitions between different states and were considered in the multiple-peak fitting of smFRET histograms.

First, we found that in the absence of any ligands or external stimuli, the GAIN domain was dynamic as shown from the three sensors (Fig. 4d, Supplementary Fig. 7). This is consistent with the overall conclusion from our cryo-EM analysis. Interestingly, we found that the GAIN domain undergoes transition between defined conformational states with respect to the 7TM domain rather than a continuum of states. Importantly, quantitative analysis revealed that in each of the three sensors transition happened between up to three distinct conformations of the GAIN domain with respect to the 7TM domain (Fig.4d-f). For the Y795:A871 sensor we found that 44% of traces showed a single FRET state, 48% showed two FRET states, and 8% showed three FRET states before photobleaching (Supplementary Fig. 8). For this sensor the three FRET states that were consistently visited were 0.44, 0.55, and 0.65 (Fig 4e, f). This is in qualitative agreement with the approximate distances between these residues in the GAIN/7TM models that were fit into the three cryo-EM conformations which are ∼61 Å, ∼57 Å, and ∼54 Å (measured from the α-carbon).

### Synthetic antibodies can activate ADGRL3 and alter the conformational distribution of GAIN/7TM

Binding of synthetic ligands to the ECR of ADGRL3 and other aGPCRs can alter receptor signaling^28,35,36,68,69^. We tested the effect of sABs LK3 and LK1 binding on ADGRL3 signaling activity using an SRE-luciferase assay^24^. LK3, which was used for cryo-EM studies, did not change the basal activity of the receptor when compared to the control experiment, acting as a neutral synthetic binder for ADGRL3 in the SRE assay (Fig. 5a). On the other hand, addition of LK1 resulted in a substantial increase in signaling compared to untreated sample or LK3 (Fig. 5a).

**Fig. 5:**
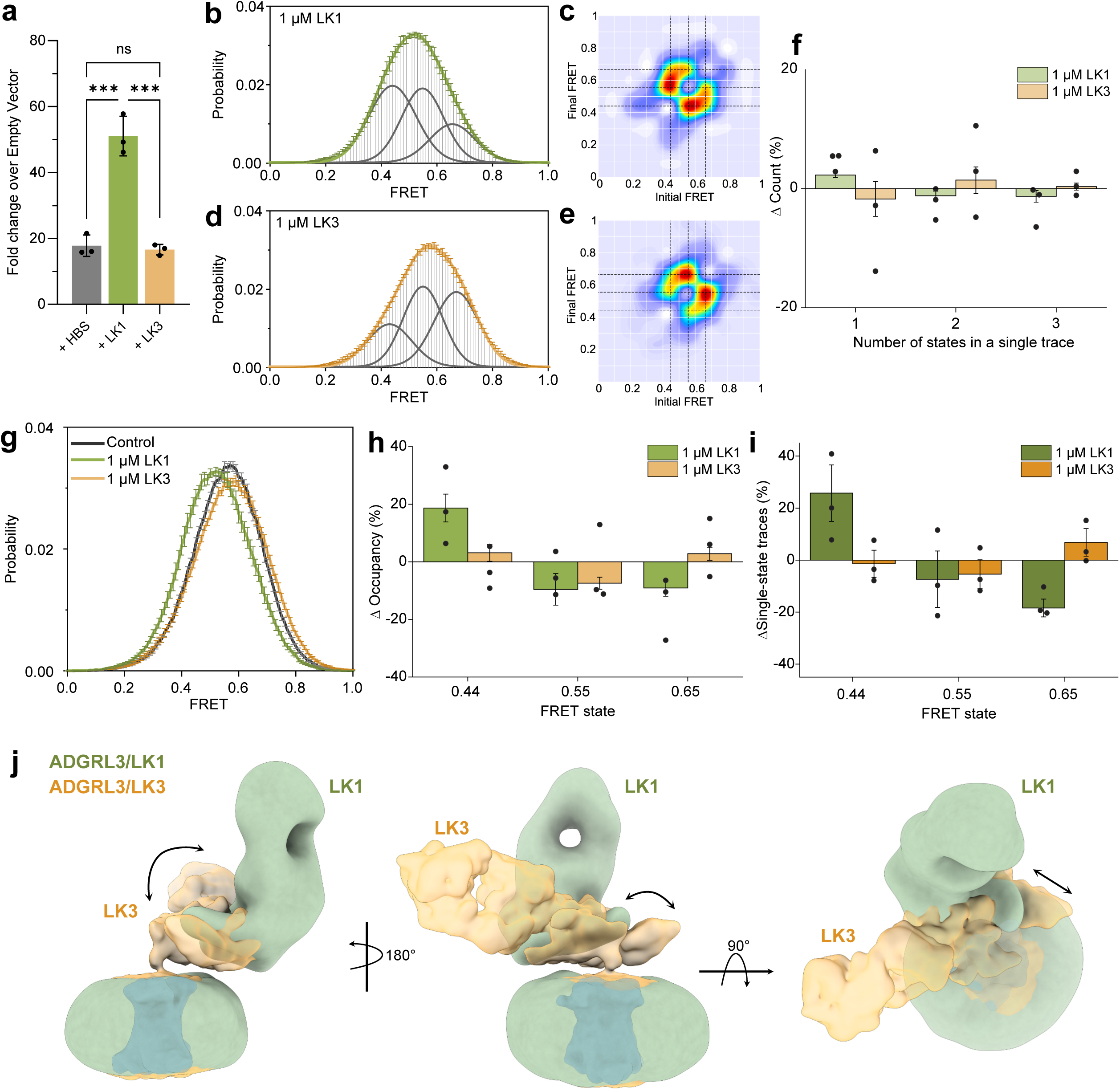
Binding of the activating sAB LK1 increases the occupancy of the lowest FRET peak. **a** SRE-luciferase assay for signaling of ADGRL3 in the presence of 1 µM purified sABs LK1 and LK3 presented as fold increase over empty vector. Data are presented as mean ± SD of three repeats (n = 3) for a representative of three independent experiments. ***p = 0.0001; vs. HBS buffer treatment; two-way ANOVA. **b** smFRET population histogram of Y795:A871 UAA sensor in the presence of 1 µM LK1 antibody. Three single Gaussian distributions (gray) were fitted to the histogram centered at 0.44, 0.55, and 0.65 with cumulative fit in green. **c** Transition density plots of Y795:A871 UAA sensor with 1 µM LK1. Dashed lines indicate the most frequent transitions. **d** smFRET population histogram of Y795:A871 UAA sensor pair in presence of 1 µM LK3 antibody. Three single Gaussian distributions (gray) were fitted to the histogram centered at 0.44, 0.55, and 0.65. Orange line represents the cumulative fit. **e** Transition density plots of Y795:A871 UAA sensor pair with 1 µM LK3. Dashed lines indicate the most frequent transitions. **f** percent change in the number of states occurring in individual traces in the presence of antibodies LK1 or LK3 versus the apo receptor. **g** smFRET population histograms of Y795:A871 UAA sensor pair in control conditions in the presence of 1 µM LK1 or 1 µM LK3. **h** Change in occupancy of the three FRET states after addition of LK1 or LK3 compared to the apo state. **i** Change in the occurrence of different FRET states among traces that showed only one state during recording between apo and LK1 or LK3 condition. **j** Superimposition of low-resolution maps of ADGRL3 in complex with sABs LK3 (in yellow) and LK1 (in green) showing shift in the HormR/GAIN domains position in relation to the micelle. (**b-i**) Data represent mean ± s.e.m., *n* = 3 from three independent biological replicates.

Next, we wondered whether sABs that affect ADGRL3 downstream signaling would alter the ECR-to-7TM orientation. We speculated that if the antibodies alter receptor signaling purely via the TA-activation model, the ECR-TM orientation would be independent of receptor signaling; whereas if the antibodies alter receptor signaling via the ECR-mediated mechanism, receptor signaling would depend on the ECR-TM orientation. To test this, we performed similar smFRET measurements with the sensor Y795UAA:A871UAA in the presence of LK1 – as the activating sAB, and LK3 – as the neutral binder. Both in the presence of 1 µM LK1 and 1 µM LK3, the ECR still remained dynamic while transitioning between the same three states as the apo receptor (Fig. 5b-e). While the percentage of traces showing one, two or three FRET states before photobleaching were the same with LK1 compared to the apo receptor (Fig. 5f), we found that the overall FRET histogram shifted towards lower FRET (Fig. 5g), increasing occupancy of the lowest FRET peak at 0.44 by 18% while decreasing the occupancy of the FRET states at 0.55 and 0.65 by about 9% in each case (Fig. 5h). Consistently, among the traces that were stable in a single FRET state before photobleaching, the percentage of traces that were in FRET 0.44 increased by 22%, but decreased for the higher FRET states (Fig. 5i). Together these results support the notion that the agonist LK1 increases the stability of the lowest FRET state. Therefore, the lowest FRET state of 0.44 is likely associated with higher receptor activity.

Unlike LK1, neutral binder LK3 did not change the occupancy of the FRET states compared to the apo receptor (Fig. 5g-i). Finally, control experiments with a non-specific sAB showed no change in FRET histogram (Supplementary Fig. 9). Taken together, these results suggest that different conformations of the ECR have different signaling capacities and specific antibodies can modulate the conformational distribution of the ECR to modulate signaling output without locking the ECR into a specific conformation.

We then used cryo-EM single particle analysis to visually observe the effect of LK1 on the ECR-7TM orientation of ADGRL3. The low-resolution cryo-EM map showed that LK1 binds to a different epitope on the GAIN domain when compared to LK3-bound model (Fig. 5j). LK1 positions itself on top of the GAIN domain, possibly pushing it down. This causes the GAIN domain to bend closer to the membrane, while moving sideways away from the central axis of 7TM region. This orientation of the GAIN domain suggests that LK1 might activate the receptor by pushing/pulling forces extorted to the ECR.

### Cancer-associated mutations at the GAIN-TM interface shift ECR-TM conformation

To further explore and confirm the relationship between the ECR conformation and receptor function we performed experiments with previously described cancer-associated mutations S810L/E811Q. These residues are positioned on one of the exposed loops of the GAIN domain that faces the 7TM in our cryo-EM model (Fig. 6a)^19,70^. Recent work showed that ADGRL3 signaling through Gα_12/13_ was impaired by the single-point cancer-related mutations S810L and E811Q^70^ although the mutations had no effect on autoproteolysis and are not a part of the TA^19^. These observations suggested that GAIN domain could make transient interactions with the extracellular loops on the 7TM.

**Fig. 6:**
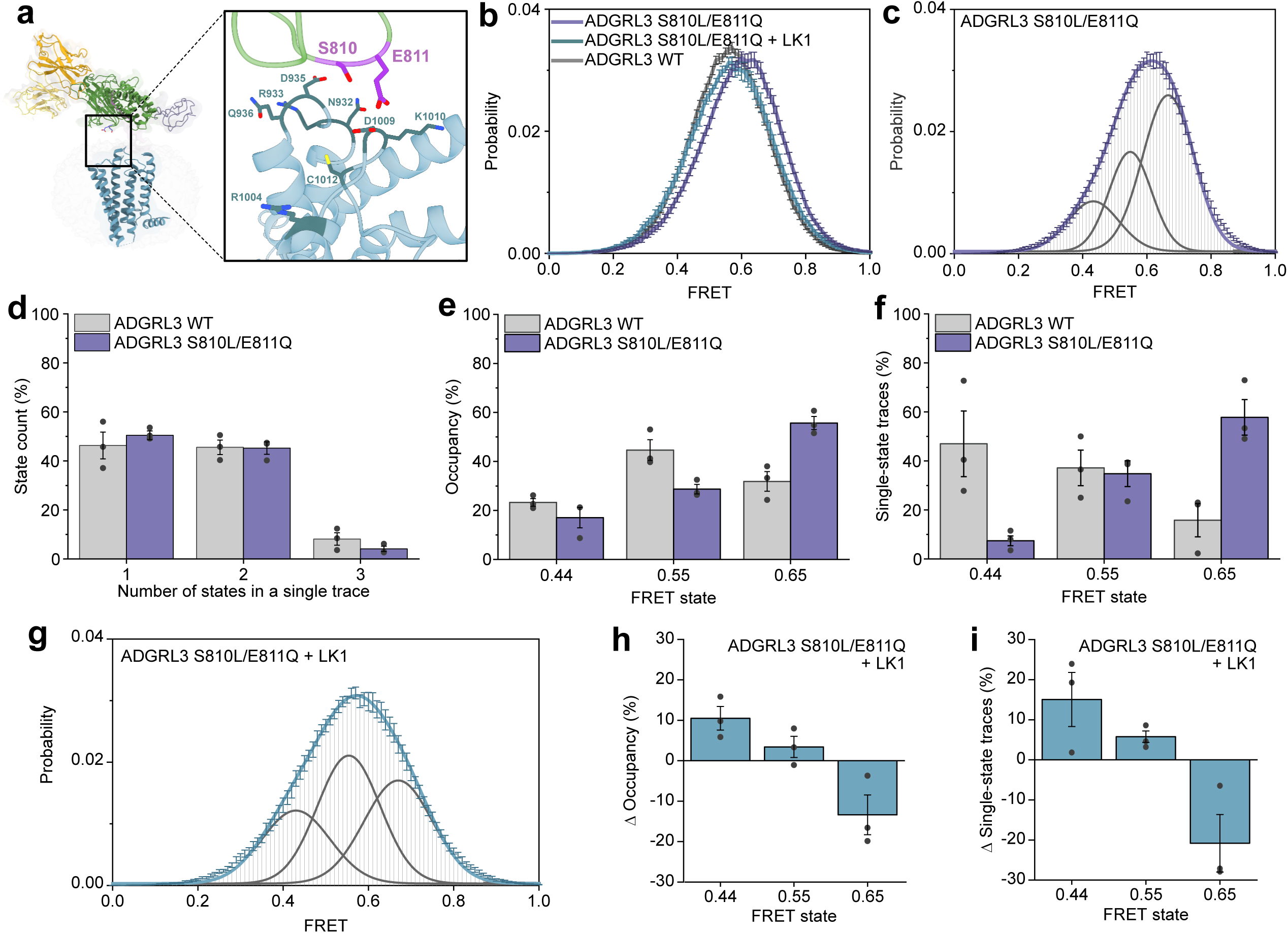
Cancer-associated GAIN domain mutant shifts the smFRET peaks to higher FRET. **a** Model of ADGRL3 showing positions of residues S810/E811 in purple, with potential interaction residues from extracellular loops of 7TM shown as sticks (blue). **b** Compilation of smFRET population histograms of ADGRL3 for the sensor Y795:A871 UAA and for mutant S810L/E811Q with or without 1 µM LK1 and ADGRL3 WT. **c** smFRET population histogram of ADGRL3 mutant S810L/E811Q fitted to three Gaussian distributions (gray) centered at 0.44, 0.55, and 0.65. The cumulative fit is purple. **d** Number of states (%) occurring in individual traces from WT ADGRL3 and S810L/ E811Q mutant. **e** Occupancy of the three FRET states in WT and S810L/E811Q mutant ADGRL3. **f** Occurrence of the three individual FRET states (0.44, 0.55, and 0.65) among single-state traces. **d - f** Data represent mean ± s.e.m. (*n = 3*), of three independent biological replicates *p < 0.05, **p < 0.01 **g** smFRET population histogram of ADGRL3 mutant S810L/E811Q in a presence of 1 µM LK1 antibody. Histogram was fitted to three Gaussian distributions (gray) with centers at 0.44, 0.55, and 0.65. Blue line is the cumulative fit. **h** Change in the occupancy of the three FRET states after addition of 1 µM LK1 to ADGRL3 mutant S810L/E811Q **i** Change in the percentage of different FRET states among all traces that stayed in a single state throughout the recording in the presence of 1 µM LK1 compared to apo receptor.

Interestingly, we found that in the presence of the S810L/E811Q mutation, the smFRET distribution shifted towards higher FRET values compared to the WT receptor (Fig. 6b, c). Quantification of FRET traces showed that there was no significant difference in the overall distribution of number of states per individual trace between the WT and mutant receptor (Figure 6d). However, the occupancy of high FRET state at 0.65 was significantly increased from 32% to 56% for the S810L/E811Q mutant compared to the WT receptor and the occupancy of other FRET states at 0.44 and 0.55 decreased (Fig. 6e). Consistent with this, among the traces that showed a single FRET state during the recording, the fraction of traces at FRET 0.65 increased from 16% to 58%, almost entirely at the expense of reduction in traces at 0.44 (Fig. 6f). Finally, we found that addition of 1 µM LK1 to this mutant receptor shifted the FRET histogram towards to the lower FRET values and similar to the WT receptor (Fig. 6g) by reducing the occupancy of the highest FRET state at 0.65 and increasing the occupancy of the lowest FRET state at 0.44 (Figure 6h, i). Together these results are consistent with the notion that the highest FRET state at 0.65 for this sensor corresponds to lower signaling propensity.

## DISCUSSION

In this work, we aimed to elucidate the structural features and conformational dynamics of ECR-bound aGPCRs to better understand the molecular nature of crosstalk between ECR and 7TM regions of aGPCRs. aGPCRs are cell-surface receptors that have key roles in a vast variety of physiological functions that range from embryogenesis to immunology and neurobiology to organogenesis. aGPCR have multidomain ECRs that participate in cell-cell adhesion, communication, and signal transduction^13,18,71^. Despite the recent emergence of structural and functional data, the conformational landscape between the ECR and 7TM regions in a holoreceptor is uncharacterized, and how exactly aGPCRs convert ECR-mediated cellular adhesion into transmembrane signaling is not known. The TA mediated activation mechanism of aGPCRs has been extensively studied and structures of TA bound transmembrane helices of aGPCRs have been determined^39–42,49^. However, multiple lines of evidence from different labs suggest that the TA-mediated mechanism may not be the only one underlying many physiological functions of aGPCRs. The cleaved TA is tightly locked in a hydrophobic pocket within the GAIN domain surrounded by 15 backbone hydrogen bonds. Based on GAIN domain structures, the force that is required to expose the TA has been reported to be as high as 200 pN^72–74^, raising the question of whether all aGPCRs detect such high physical forces on the receptor in a cellular environment. This result implies that aGPCRs are insensitive to forces in the range of 2-40 pN where most receptor-ligand mediated biological forces fall^75–78^ or to compression forces that push on the receptor which may be common during synapse formation and embryogenesis^79–81^ In addition, some aGPCRs such as ADGRLs and CELSRs function independent of autoproteolysis, which is a prerequisite for TA-dependent mechanism^7,29,31^

### The GAIN domain has stable positions with respect to the TM domain

Here we report the overall architecture of the ECR-bound ADGRL3 and define the relative position of the GAIN domain to the 7TM and membrane. We found that the GAIN domain lays flat, in close proximity to the membrane and does not display a large extent of flexibility, limited to 45⁰ range of movement. 3D variability tools showed that GAIN samples other conformations, all of which maintain the overall arrangement of the domains. Furthermore, smFRET experiments with full-length ADGRL3 show that the GAIN domain adopts three stable conformations with respect to the 7TM domain. These conformations have slow exchange rates between each other and switch from one to another in an orderly manner. With these results, we propose that the GAIN/7TM of ADGRL3 displays a restricted flexibility, with a limited range of motion between the two regions of the receptor, and adopting distinct and stable conformations (Fig. 7a). Though our interpretation seems contrary to previous studies that reported high flexibility of the ECR, it is possible that ECR stays more rigid in the basal state and becomes more flexible when the receptor is activated, as numerous such studies captured the active conformation of the 7TM complexed with trimeric G proteins.

**Fig. 7:**
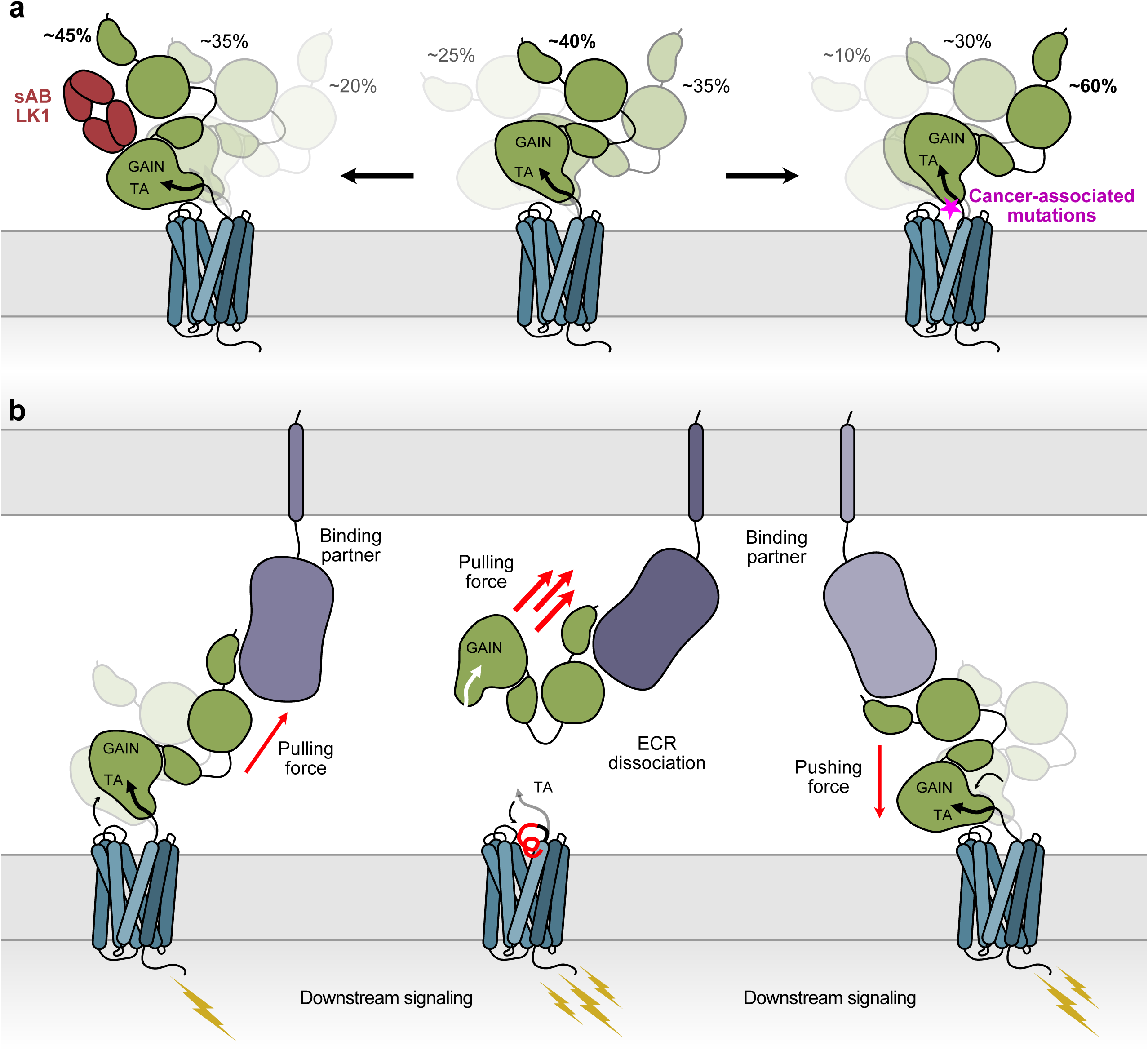
Antibodies and disease mutations at the ECR/7TM interface change the distributions of ADGRL3 holoreceptor conformations. **a** Schematic cartoon of the effects of sAB LK1 and cancer-associated mutations on conformational dynamics of ECR in ADGRL3. Activating sAB LK1 stabilizes the lower FRET states, while cancer mutants stabilize the higher FRET states of ADGRL3. **b** Proposed ECR-mediated activation mechanisms of aGPCRs. Binding of an extracellular protein ligands subject aGPCR to range of potential forces. Small pulling force can activate the holoreceptor by changing conformation of the ECR and inducing the signal trandsuction by 7TM region. Greater pulling forces can lead to dissociation of the ECR which triggers the TA activation mechanism. Binding partners can also push on the ECR and activate the receptor by shifting the ECR conformation.

Importantly, we have not observed the TA peptide dissociation from the GAIN domain or partial exposure to the 7TM domain. The high-resolution map of the GAIN domain from ECR-bound ADGRL3 includes the density of the intact TA peptide, suggesting that the TA peptide, especially the conserved F857, L860 and M861 residues, that dive deep into the 7TM domain to enable TA-mediated activation, are not exposed to the 7TM when the receptor is at its basal state.

Interestingly, the orientation of GAIN/HormR fragment in the model positions the HormR domain in close proximity to the membrane and 7TM region. This suggests that other more N-terminal domains of the ECR can potentially also remain closer to the membrane, forming a compact multidomain module, rather than freely sampling extracellular space in search of binding partners. The structures of isolated full ECR of both ADGRG6/GPR126^32^ and ADGRC1/CELSR1^82^ present compact conformations; similarly, an AlphaFold2 prediction of ADGRL3 full-length ECR places the Lec/Olf domains close on top of the extracellular facing surface of the GAIN domain, making a compact module (Supplementary Fig. 10). Furthermore, several functional studies have shown that deletion of N-terminal domains from different aGPCRs including ADGRG1/GPR56, ADGRG6 and ADGRL1 affected the basal activity of the receptors^24,32,34^. In addition, the same conclusion was achieved from the observations that the binding of antibodies or endogenous ligands to the most N-terminal domains of aGPCRs also changed receptor signaling and function^28,35,36,68,69^. Put together, this suggests that the ECR of some aGPCRs may adopt a tightly packed conformation that transmits information from the N-terminal to C-terminal ECR domains, and to the 7TM domain.

### Agonistic antibodies and disease mutations at the ECR-TM interface change the ECR-TM conformation

LK1 antibody which binds to the ECR and modulates receptor activity, also changes the distribution of the population of conformations in the smFRET assays indicating a correlation between each ECR conformation and the signaling level of the receptor (Fig. 7a). This implies that the activation by the antibody does not require ECR dissociation as no dissociation was observed in smFRET traces. A low-resolution model of LK1 bound to ADGRL3 further confirms the intact LK1/ADGRL3 complex and presents a different orientation of the GAIN domain. Similarly, cancer-associated mutations that are shown to decrease the receptor activity also change the population of smFRET conformations, suggesting that a shift in the populations of conformations might underly reduction in receptor signaling^70^ (Fig. 7a). . A local change in the transient interdomain interaction of the GAIN and 7TM domains likely induces small and tunable changes in receptor activity. Our data suggest that the highest FRET state corresponds to the lower receptor signaling and the lowest FRET state corresponds to the higher basal receptor signaling

### aGPCRs may use an ECR-mediated activation mechanism to sense pushing as well as pulling forces

Many biological forces are smaller than 200 pN, the force that is needed to separate the TA from the GAIN domain^75–78^. To sense these smaller forces, and to regulate aGPCR function on and off, a mechanism that does not depend on ECR dissociation and TA exposure might be at work. At low force or no force conditions, aGPCR may be reversibly regulated by binding and dissociation of a ligand to the ECR without ECR shedding and TA exposure (Fig. 7b). In this model, the ECR-7TM communication is altered by transient interactions between ECR and 7TM. The TA peptide remains at its original position and is not involved in signaling. Because the TA-mediated mechanism is a “one and done” mechanism that is irreversible and prevents the receptor from going back to its inactive resting state, the ECR-mediated mechanism may operate in situations where a reversible regulation is needed. The ECR-mediated mechanism may also enable responding to compressing forces on the receptor, that directly “push” on the protein (Fig. 7b). In cases where a large “pulling” force is executed on the ECR, the ECR may be removed from the 7TM releasing the tethered agonist and activating the aGPCR irreversibly but acutely (Fig. 7b). Future work that dissects different activation mechanisms of aGPCRs in different physiological contexts will shed light on this fascinating family of receptors.

## METHODS

### Protein expression and purification

ADGRL3 protein constructs were expressed using the baculovirus method. *Spodoptera frugiperda* Sf9 cells were transfected with specific plasmid and linearized baculovirus DNA (Expression Systems, 91-002) using Cellfectin II (Thermo Fisher, 10362100). Then, baculovirus was amplified in Sf9 cells in SF-900 III medium supplemented with 10% FBS (Sigma–Aldrich, F0926).

Construct containing Avi-tagged HormR/GAIN domains of ADGRL3 was expressed in High Five cells (Thermo Fisher, B85502) grown in Insect-Xpress medium (Lonza, 12-730Q), as described previously^35^. Briefly, cultures were infected with HormR/GAIN baculovirus at a density of 2.0 × 10^6^ cells ml^−1^ and incubated for 72 h at 27 °C. The cells were pelleted by centrifugation and the conditioned medium containing the secreted glycosylated proteins were collected. Final concentrations of 50 mM Tris pH 8, 5 mM CaCl_2_, and 1 mM NiCl_2_ were added to the media and stirred for 30 min. Formed precipitation was then removed by centrifugation at 8000*g* for 30 min. The clear supernatant was later incubated with Ni-NTA resin (Qiagen, 30250) for 3 h. Nickel beads were collected using a glass Buchner funnel connected to a vacuum trap and washed with 10 mM HEPES pH 7.2, 150 mM NaCl, 20 mM imidazole. HormR/GAIN domains were then biotinylated on-column with 50 mM bicine pH 8.3, 10 mM MgOAc, 100 mM NaCl, 10 mM ATP, 0.5 mM biotin and 5mM BirA at 27°C with gentle mixing. The protein was eluted from the resin with 10 mM HEPES pH 7.2, 150 mM NaCl, 200 mM imidazole, and the concentrated elution was injected onto Superdex S200 10/300 GL column (GE Healthcare) equilibrated with 10 mM HEPES; pH 7.2, 150 mM NaCl. Purified fractions of the complex were used for further experiments.

The HormR+GAIN/7TM construct of ADGRL3 was expressed in Sf9 insect cells grown in ESF 921 serum-free media (Expression System). Cells grown to a density of 4.0 × 10^6^ cells ml^−1^ were infected with high-titer baculovirus. Cultures were grown at 27 °C and harvested by centrifugation 50h post infection. Cells were lysed in 10mM Tris pH 7.5, 1mM EDTA, 2 mg ml^−1^ iodoacetamide supplemented with cOmplete Protease Inhibitor Cocktail tablets (Roche) in Dounce homogenizer. Membrane fractions were collected by centrifugation at 30,000*g* at 4 °C for 1 h, and solubilized in buffer containing 30 mM HEPES pH 7.5, 500 mM NaCl, 10% glycerol, 1% (w/v) n-Decyl-β-D-Maltopyranoside (DM, Anatrace), 0.2% (w/v) cholesteryl hemisuccinate (CHS), 2 mg ml^−1^ iodoacetamide supplemented with Complete Protease Inhibitor Cocktail tablets (Roche) for 1 h at 4 °C. Cellular debris was removed by centrifugation at 100,000*g* for 1 hour and the solubilized proteins were purified from the supernatant over M1 anti-Flag affinity resin in the presence of 2 mM CaCl_2_. The resin was washed with 10 column volumes of 30 mM HEPES pH 7.5, 150 mM NaCl, 0.01% lauryl maltose neopentyl glycol (LMNG, Anantrace), 0.005% (w/v) CHS, 2 mM CaCl_2_ before bound material was eluted in buffer containing 30 mM HEPES pH 7.5, 150 mM NaCl, 0.01% LMNG, 0.0025% (w/v) GDN, 8 mM EDTA and 0.2 mg ml^−1^ Flag peptide. The purified receptor was subsequently concentrated using an Amicon Ultra Centrifugal Filter (MWCO 100 kDa) and run on size-exclusion chromatography on a Superdex 200 10/300 column (GE Healthcare) pre-equilibrated with 30 mM HEPES pH 7.5, 150 mM NaCl, 0.00075% (w/v) LMNG, 0.00025% (w/v) GDN, 0.001% (w/v) CHS. Eluted fractions of the receptor were pooled and concentrated. Final yield of purified ADGRL3 was approximately 2.5 mg l^−1^ of insect cell culture.

### Negative-stain electron microscopy analysis

Purified receptor was diluted to ∼5 ug mL^−1^ and applied to freshly glow-discharged EM carbon coated copper grid (Electron Microscopy Sciences, CF400-Cu,) for 30 sec. The protein was blotted off with filter paper (Sigma–Aldrich, WHA1001110), and immediately after blotting, the grid was then touched to a 25 μL drop of 1% uranyl formate solution for 30 sec and blotted off, followed by air dry. The negatively-stained sample was imaged at RT with a Tecnai G2 F30 operated at 300 kV. Images were recorded at a magnification of 49 000x and processed using EMAN2 software^83^.

### Phage Display Selection

Phage Display Selection for HormR/GAIN fragment of ADGRL3 was performed according to previously published protocols^56,84^. For the first round of selection 200 nM of protein was immobilized on streptavidin magnetic beads. The beads were washed three times to remove unbound target protein. Next, to prevent nonspecific binding of the phage, 5 mM D-biotin was added to block unoccupied streptavidin on the beads. Then, the beads were incubated for 30 min at RT with the phage library E^85^, containing 10^12^-10^13^ virions ml^-1^ with gentle shaking and washed to remove unbound phages. Beads with bound phages were used directly to infect log phase *E. coli* XL1-Blue cells. Cells were grown overnight in 2YT media with 50 µg/mL ampicillin and 10^9^ p.f.u. ml^-1^ of M13 KO7 helper phage in order to amplify phages. Three additional rounds of selection were performed with decreasing target concentration in each round (100 nM, 50 nM, 10 nM) using the amplified pool of virions of the prior round used as the input. Those rounds were performed using semi-automated system with the Kingfisher instrument. In rounds 2 to 4 phages were eluted using 0.1 M glycine pH 2.7.

### Enzyme-Linked Immunoabsorbent Assays (ELISA)

ELISA experiments were carried out using a 96-well flat-bottom plate coated with 50 µL of 2 mg ml^-1^ neutravidin in Na_2_CO_3_ pH 9.6 and subsequently blocked with 0.5% Bovine Serum Albumin in 1×PBS. Binding screens of all of the selected sABs in phage format was performed using a single point phage ELISA. 400 µL of 2YT media with 100 µg ml^-1^ ampicillin and M13 KO7 helper phage were inoculated with single *E. coli* XL1-Blue colonies harboring phagemids, and cultures were grown at 37°C for 18h in a 96-deep-well block plate. The cells were pelleted by centrifugation and sAB phage-containing supernatants were diluted 20× in ELISA buffer. Diluted phages were then applied to ELISA plates, preincubated for 15 min with 50 nM of biotinylated target proteins at RT. Plates with added phages were incubated for 15 min at RT and washed 3-times with 1×PBST. The washing step was followed by 30 min incubation with HRP-conjugated Mouse Anti-M13 monoclonal antibody (GE Healthcare, 27942101) diluted in PBST in 1:5000 ratio. Excess antibody was washed away with 1×PBST and plates were developed using TMB substrate, quenched with 1.0 M HCl and the signal was determined by absorbance measurement (A_450_).

### Cloning, Overexpression and Purification of sABs

Phage ELISA results were used to select sAB clones that were sequenced at DNA Sequencing Facility at The University of Chicago. In-fusion cloning^86^ was used to reformat unique sABs clones into pRH2.2, an IPTG inducible vector for bacterial expression.

*E*. *coli* BL21 (Gold) cells were transformed with sequence-verified sAB plasmids. Cultures were grown in 2YT media supplemented with 100 μg mL^-1^ at 37°C until they reach OD_600_ = 0.8, when they were induced with 1 mM IPTG. The culture was continued for 4.5h at 37°C and cells were harvested by centrifugation. Cell pellets were resuspended in 20 mM HEPES pH = 7.5, 200 mM NaCl, 1 mM PMSF, 1 μg ml^-1^ DNase I, and lysed by ultrasonication. The cell lysate was incubated at 60°C for 30 min. Heat-treated lysate was centrifuged at 50,000 *x g* to remove cellular debris, filtered through a 0.22 μm filter and loaded onto a HiTrap protein L (GE Healtchcare) column pre-equilibrated with 20 mM HEPES pH 7.5 and 500 mM NaCl. The column was washed with 20 mM HEPES pH 7.5 and 500 mM NaCl and sABs were eluted with 0.1 M acetic acid. Protein containing fractions were loaded directly onto an ion-exchange Resource S column pre-equilibrated with 50 mM NaOAc pH 5.0 and washed with the equilibration buffer. sABs elution was performed with a linear gradient 0–50% of 50 mM NaOAc pH 5.0 with 2 M NaCl. Purified sABs were dialyzed overnight against 20 mM HEPES pH 7.5 with 150 mM NaCl. The quality of purified sABs was analyzed by SDS-PAGE.

### Formation of receptor/sAB Complex

Receptor/sAB complex was formed by mixing 1.5-fold molar excess of the sAB with the receptor and 30 min incubation on ice. The complex was subjected SEC on a Superdex 200 10/300 column pre-equilibrated with 30 mM HEPES pH 7.5, 150 mM NaCl, 0.00075% (w/v) LMNG, 0.00025% (w/v) GDN, 0.001% (w/v) CHS. Formation of the complex was determined by retention volume analysis of the complex with respect to that of target alone and co-elution of the individual components on SDS-PAGE.

### Cryo-EM data acquisition

For cryo-EM analysis, the purified 7TM ADGRL3/sAB complex was concentrated to 2.5 mg ml^−1^. Right before sample processing, fluorinated octyl maltoside (FOM) was added to the solution to final concentration of 0.05%. Vitrified specimen was prepared by applying 3 μl of protein complexes onto a glow-discharged 300 mesh gold holey carbon Quantifoil R 1.2/1.3 grid (Quantifoil Micro Tools) and frozen in liquid ethane cooled by liquid nitrogen inside a Vitrobot Mark IV (FEI). Cryo-EM imaging was performed using Titan Krios electron microscope operated at 300 kV (Thermo Scientific) equipped with a K3 direct electron detector (Gatan, Inc.). Movies were recorded with a nominal magnification of ×64,000 in super-resolution counting mode, corresponding to a pixel size of 0.67 Å on the specimen level. 6976 movies were recorded with defocus values in the range of −1.0 to −2.0 μm, using an accumulated dose rate of 65 electrons per Å^2^ and a total of 58 frames per movie stack.

### Cryo-EM data processing and 3D reconstructions

Movies were imported to CryoSPARC 4.4^61^ and stack images were subjected to full-frame and local motion correction, as well as contrast transfer function (CTF) estimation. In total, 2,621,532 particles were selected using automated particle picking. Multiple rounds of reference-free 2D classification were performed in order to discard particles grouped in poorly defined classes, resulting in 549,473 particles with well-defined features for the detergent micelle, ECR and sAB for further processing. 3D Ab-initio reconstruction (with 3 classes) generated the initial reference maps and Heterogeneous Refinement, followed by Non-Uniform Refinement was used to further classify particles and refine structures. 242,439 particles from the best 3D map were then used for 3D Variability analysis^60^, and separated frames from this analysis were subjected to two rounds of Heterogenous Refinement. Then, classes were subjected to iterative rounds of Non-Uniform Refinement, with the best class accommodating 96,958 particles generating a map with an indicated global resolution of 5.5 Å. To further improve the quality of the extracellular domains in the structure, the 5.5 Å map of the entire complex was used in Local Refinement job, utilizing a mask focusing on the HormR+GAIN+sAB fragment. Multiple rounds of Non-Uniform and Local Refinements produced a higher resolution map with a global resolution of 3.9 Å. The detailed data processing flow is shown in Supplementary Fig. 2. Reported resolutions are based on the gold-standard Fourier shell correlation (FSC) using the 0.143 criterion. Local resolution was determined using half-reconstructions as input maps.

### Model building and refinement

The AlphaFold2 predictions of HormR/GAIN of ADGRL3 and sAB LK3 were used for model building^59^. Models were manually fitted into the density map of HormR+GAIN/LK3 using the Fit in Map function of UCSF ChimeraX^87^. Then, the model was docked into the EM density map using Phenix dock in map function, refined using the real-space refinement module in the Phenix software suite^88^ and then manually checked and adjusted residue-by-residue to fit the density using COOT^89^, in an iterative manner. Structural figures were created using UCSF ChimeraX^87^.

### Serum Response Element Luciferase Assays

HEK293T cells (ATCC, CRL-3216) were seeded on a 96-well flat-bottom plate precoated with 0.5% gelatin and grown until 50-60% confluent in DMEM supplemented with 10% (v/v) FBS. Cells were then co-transfected with full-length ADGRL3 (1 ng well^-1^)^24,35^, Dual-Glo luciferase reporter plasmid (20 ng well^-1^)^34^, using 0.3 μL LipoD293T (SL100668; SignaGen Laboratories). DNA levels were balanced among transfections by addition of the empty pCMV5 vector to 100 ng total DNA. Eighteen hours after transfection media was aspirated and replaced with DMEM without FBS. When sABs were tested, 1 µM of sAB was added 5h after start of the serum-starvation. After 10h of serum starvation cells were lysed using Dual-Glo Luciferase Assay System from Promega and firefly and renilla luciferase signals were measured using a Synergy HTX (BioTek) luminescence plate reader. The firefly:renilla ratio for each well was calculated and normalized to empty vector. Data were then analyzed using GraphPad Prism 9.3.1 software and presented as mean ± SD of three repeats (n = 3) for a representative of three independent experiments

### Molecular cloning of smFRET constructs

The C-terminal eGFP-tagged human ADGRL3 construct was cloned in pcDNA3.1(+) vector with 12 amino acids linker and verified by sequencing (ACGT Inc). Full length ADGRL3 constructs with an amber codon (TAG) mutation of amino acid Y795 and A871, R933 or E1099 as well as variants harboring cancer mutations S8110L/E811Q were generated with QuickChange site-directed mutagenesis (Agilent). All constructs were verified with full-length sequencing (Primordium Labs). DNA restriction enzymes, DNA polymerase and DNA ligase were from New England Biolabs. Plasmid preparation kits were purchased from Macherey-Nagel.

### Transfection and Protein Expression for smFRET Experiments

HEK293T cells (Sigma) were maintained in DMEM (Gibco) supplemented with 10% (v/v) serum (Cytiva), 100 unit/mL penicillin-streptomycin (Gibco) and 15 mM HEPES (pH=7.4, Gibco) at 37 °C and 6% CO2. The cells were passaged with 0.05% trypsin-EDTA (Gibco). For unnatural amino acid-containing protein expression, the growth media was supplemented with 0.25 mM trans-Cyclooct-2-en – L – Lysine (SiChem, #SC-8008). All media was filtered by 0.2 µM PES filter (Fisher Scientific).

24 hours before transfection, HEK293T cells were cultured on poly-L-lysine-coated 18 mm glass coverslips (VWR) at 60% - 70% confluency. Two hours before transfection, the media was changed to the growth media supplemented with 0.25 mM trans-Cyclooct-2-en – L – Lysine ADGRL3 plasmids with amber codon and pNEU-hMbPylRS-4xU6M15 (pNEU-hMbPylRS-4xU6M15 was a gift from Irene Coin, Addgene plasmid # 105830) were co-transfected (1:1 w/w) into cells using Lipofectamine 3000 (Fisher Scientific) (total plasmid: 2 µg/18 mm coverslip). After 48 hours supplemented growth media was removed and cells were washed by extracellular buffer solution containing (in mM): 128 NaCl, 2 KCl, 2.5 CaCl2, 1.2 MgCl2, 10 sucrose, 10 HEPES, pH=7.4 and were kept in growth medium without trans-Cyclooct-2-en – L – Lysine for 30 min. Site-specific labeling with Cy3 and Cy5 fluorophores was done by diluting Cy3 and Cy5 alkyne dyes (Click Chemistry Tools) to the final concentration of 40 µM each in the extracellular buffer and incubating for 15 minutes in the incubator. After labelling, coverslips were gently washed by the extracellular buffer solution to remove excess dye. Live-cell images of labeled cells were collected using a Zeiss LSM800 confocal microscope with a Plan-Apochromat ×40 objective (Zeiss, 1.3 numerical aperture, oil immersion) and Zen Blue software.

### Single-molecule FRET measurements

Single-molecule experiments were conducted in chambers prepared from glass coverslips (VWR) and microscope slides (Fisher Scientific) passivated with mPEG (Laysan Bio) and biotin-PEG as previously described^66^. Prior to experiments, flow cells were functionalized with NeutrAvidin (Fisher Scientific) and anti-GFP antibody (Abcam, #ab6658).

After labeling, cells were incubated in DPBS without Ca2+/Mg2+ for 15 minutes and harvested. Afterwards cells were resuspended in 100 µL lysis buffer consisting of 200 mM NaCl, 50 mM HEPES, 1 mM EDTA, protease inhibitor tablet (Fisher Scientific), and 0.1 w/v% LMNG-CHS (10:1, Anatrace), pH 7.4. Cells were lysed with gentle mixing at 4 °C for 1 hour. Cell lysate was then centrifuged for 20 min at 20,000 g and 4 °C. The supernatant was collected and kept on ice. Diluted cell lysate was immobilized in the imaging chamber and imaged in imaging buffer consisting of (in mM) 128 NaCl, 2 KCl, 2.5 CaCl2, 1.2 MgCl2, 40 HEPES, 4 Trolox, 0.005 w/v% LMNG-CHS (10:1), 0.0004 w/v% GDN, and an oxygen scavenging system consisting of protocatechuic acid (Sigma) and 1.6 U/mL bacterial protocatechuate 3,4-dioxygenase (rPCO) (Oriental Yeast Co.), pH 7.35. Samples were imaged with a 100× objective (Olympus, 1.49 NA, Oil-immersion) on a custom-built microscope with 30 ms time resolution unless stated otherwise. 532 nm and 638 nm lasers (RPMC Lasers) were used for donor and acceptor excitation, respectively.

### smFRET data analysis

Analysis of single-molecule fluorescence data was performed as described before^65^. Particles that showed acceptor signal upon donor excitation with acceptor intensity greater than 10% above background, were automatically selected and donor and acceptor intensities were measured over all frames. Out of this pool (400-600 total), particles that showed a single donor and a single acceptor bleaching step during the acquisition time, showed stable total intensity (I_D_ + I_A_), anti-correlated donor and acceptor intensity behavior without blinking events, and lasted at least 4 seconds before bleaching were manually selected for further analysis (∼20-30% of total molecules per movie). A subset of the data was analyzed by two individuals independently and the results were compared and showed to be identical. FRET efficiency was calculated as (I_A_ − 0.085×I_D_) / (I_D_ + (I_A_ − 0.085×I_D_)), where I_D_ and I_A_ are raw donor and acceptor intensities, respectively. Data was collected for at least three independent biological replicates, and the results were averaged. Population smFRET histograms were generated by compiling at least 250 total FRET traces of molecules from all replicates. To ensure that each trace contributes equally, area under FRET histogram of individual molecules was normalized to one. Error bars on histograms represent the standard error of mean of data.

A hidden Markov model (HMM) using vbFRET software was used to idealize FRET traces^90,91^ and generate the transition density plots with number of states set to one to five states. TDP plots for each replicate showed three definitive FRET states. This was also corroborated with manual inspection of all the traces.

Next, peak fitting analysis on population smFRET histograms was performed with OriginPro and used three Gaussian distributions as 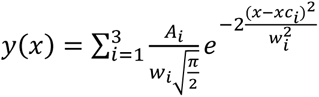, where *A* is the peak area, *w* is the peak width, and *xc* is the peak center. Peak areas were constrained to A > 0. Peak widths were constrained to 0.1 < *w* < 0.17. Peak centers were constrained to ± 0.02 of mean FRET efficiency of each FRET state obtained from the TDP plots.

## ACKNOWLEDGEMENTS

We thank Dr. James Fuller and the staff at The University of Chicago Advanced Electron Microscopy Core Facility (RRID:SCR_019198) for assistance with cryo-EM data collection. We also thank Dr Andrew Kruse and Dr Cheng Zhang for helpful discussions regarding membrane protein expression and purification, Dr. James Fuller, Dr. Jingxian Li, Dr. Navid Bavi, and Dr. Minglei Zhao for valuable advice regarding cryo-EM sample preparation and data processing. This work was supported by grants R35 GM148412 (to D.A.), R01 GM134035-01 (to D.A.), R01 GM140272 (to R.V.) F32 GM142266 (to S.J.B.), R01 GM117372 (to A.A.K.) and C-093 from the Chicago Biomedical Consortium (to D.A. and R.V.).

## DATA AVAILABILITY

The authors declare that all data supporting the findings of this study are available within the article and upon a reasonable request to the corresponding authors.

## AUTHOR CONTRIBUTIONS

S.P.K., K.C., R.V and D.A. designed all experiments and interpreted results. K.C. and K.L. cloned constructs for smFRET and protein expression and purification experiments. S.P.K. expressed and purified all proteins (with assistance from K.L.), prepared EM samples, performed specimen screening, data collection and the cryo-EM data analysis. S.J.B. assisted with cryo-EM map calculation and built and refined the HormR/GAIN structure. K.C. prepared smFRET specimens, performed smFRET measurements (with assistance from G.S.) and data analysis. P.D. carried out phage display selection and sABs characterization. S.P.K. performed cell-based signaling assays. D.A., S.P.K., R.V. and K.C. wrote the manuscript with assistance from, S.J.B. and P.D.. D.A. and R.V. supervised the project.

