## Supplementary Figures 1-10 for "Structural analysis and conformational dynamics of a holo-adhesion GPCR reveal interplay between extracellular and transmembrane domains"

for

### Structural analysis and conformational dynamics of a holo- adhesion GPCR reveal interplay between extracellular and transmembrane domains

Szymon P. Kordon[1, 2, 3, #], Kristina Cechova[4, #], Sumit J. Bandekar[1, 2, 3], Katherine Leon[1, 2, 3], Przemysław Dutka[1, 5],

Gracie Siffer[4], Anthony A. Kossiakoff[1], Reza Vafabakhsh[4, *], Demet Araç[1, 2, 3, *]

1 Department of Biochemistry and Molecular Biology, The University of Chicago, Chicago, IL, USA

2 Neuroscience Institute, The University of Chicago, Chicago, IL, USA

3 Institute for Biophysical Dynamics, University of Chicago, Chicago, IL, USA

4 Department of Molecular Biosciences, Northwestern University, Evanston, IL, USA

5 Current affiliation: Department of Structural Biology, Genentech, South San Francisco, CA, USA

### These authors contributed equally to this work

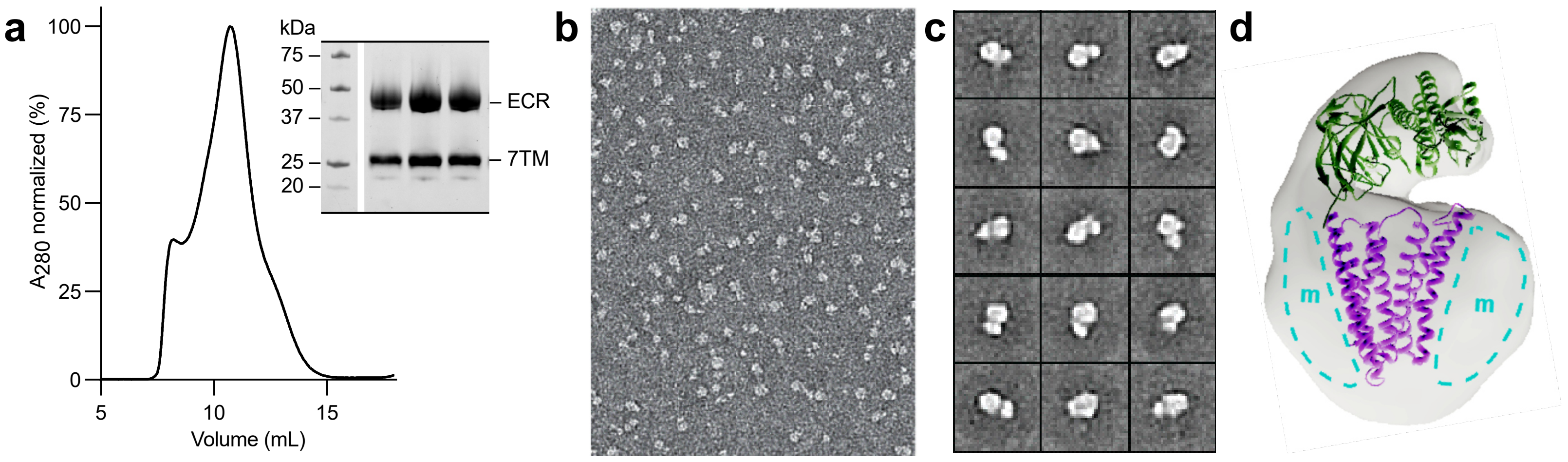


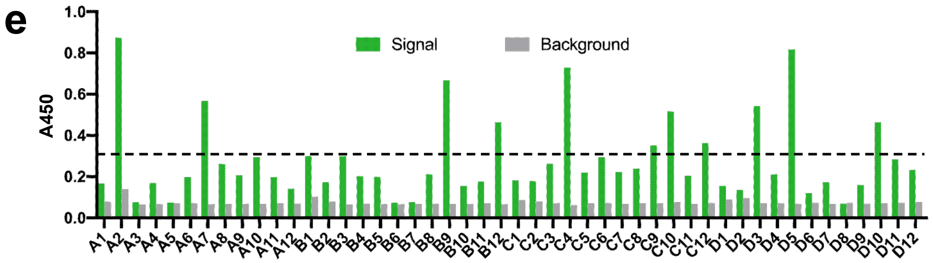


f

| sAB ID | ADGRL3 | | | | | |
|---|---|---|---|---|---|---|
| | **HormR/GAIN** | | | **Full ECR** | | |
| | $k_{on}$ ($M^{-1}s^{-1}$) | $k_{off}$ ($s^{-1}$) | $K_D$ (nM) | $k_{on}$ ($M^{-1}s^{-1}$) | $k_{off}$ ($s^{-1}$) | $K_D$ (nM) |
| **LK1** | $1.3 \times 10^5$ | $4.3 \times 10^{-4}$ | 3.3 | $1.2 \times 10^5$ | $4.8 \times 10^{-4}$ | 4.0 |
| **LK3** | $4.5 \times 10^5$ | $1.7 \times 10^{-3}$ | 3.9 | $3.5 \times 10^5$ | $1.9 \times 10^{-3}$ | 5.4 |

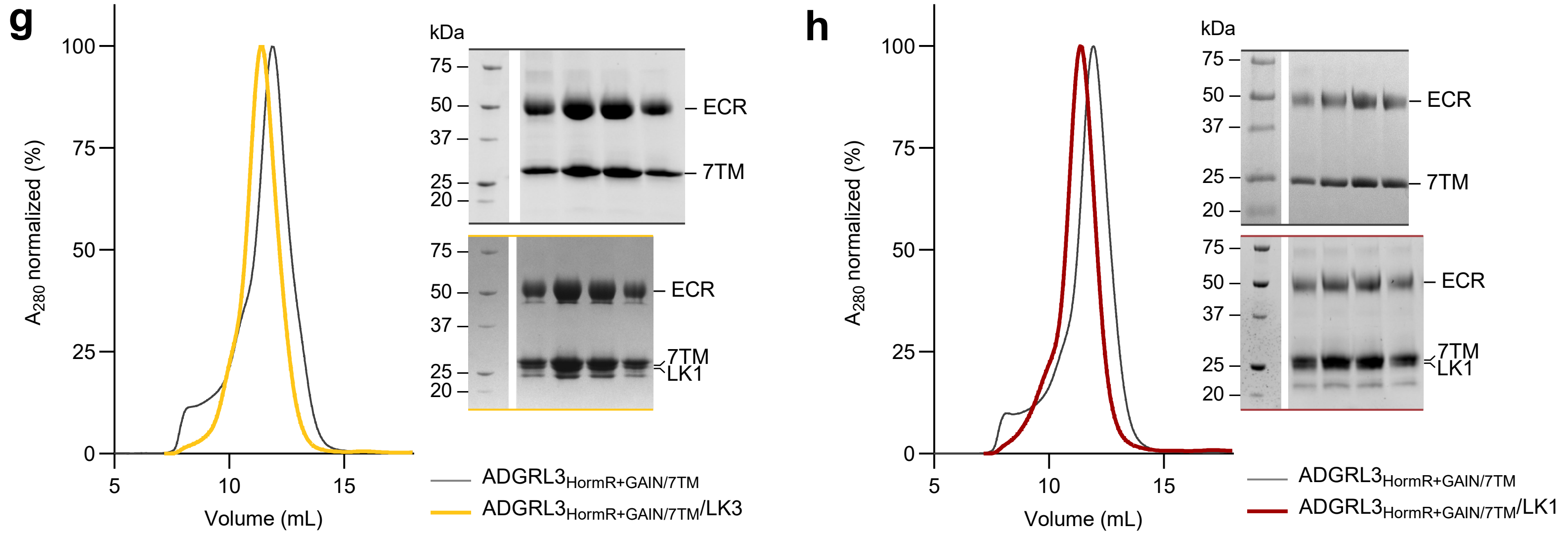


**Supplementary Fig. 1: Purification of ADGRL3$_{HormR+GAIN/7TM}$ and synthetic antigen binders generation and validation.**

**a** SEC profile and SDS-PAGE analysis of the ADGRL3 holoreceptor purified in detergent. **b** Representative negative stain EM micrograph of purified ADGRL3 holoreceptor sample. **c** Representative negative stain EM 2D class averages of purified ADGRL3 in detergent micelle. **d** Low-resolution negative stain EM map of ADGRL3 in detergent micelle. Available structures of GAIN and 7TM domains placed manually in the density. **e** Representative results of single point phage ELISA for sABs against ADGRL3 HormR/GAIN fragment. **f** Kinetic values of sABs LK1 and LK3 developed against HormR/GAIN domains of ADGRL3 obtained by SPR. **g** and **h** SEC profiles and SDS-PAGE analyses show both LK3 (g) and LK1 (h) forming monodisperse complexes with the purified ADGRL3 in detergent micelle.

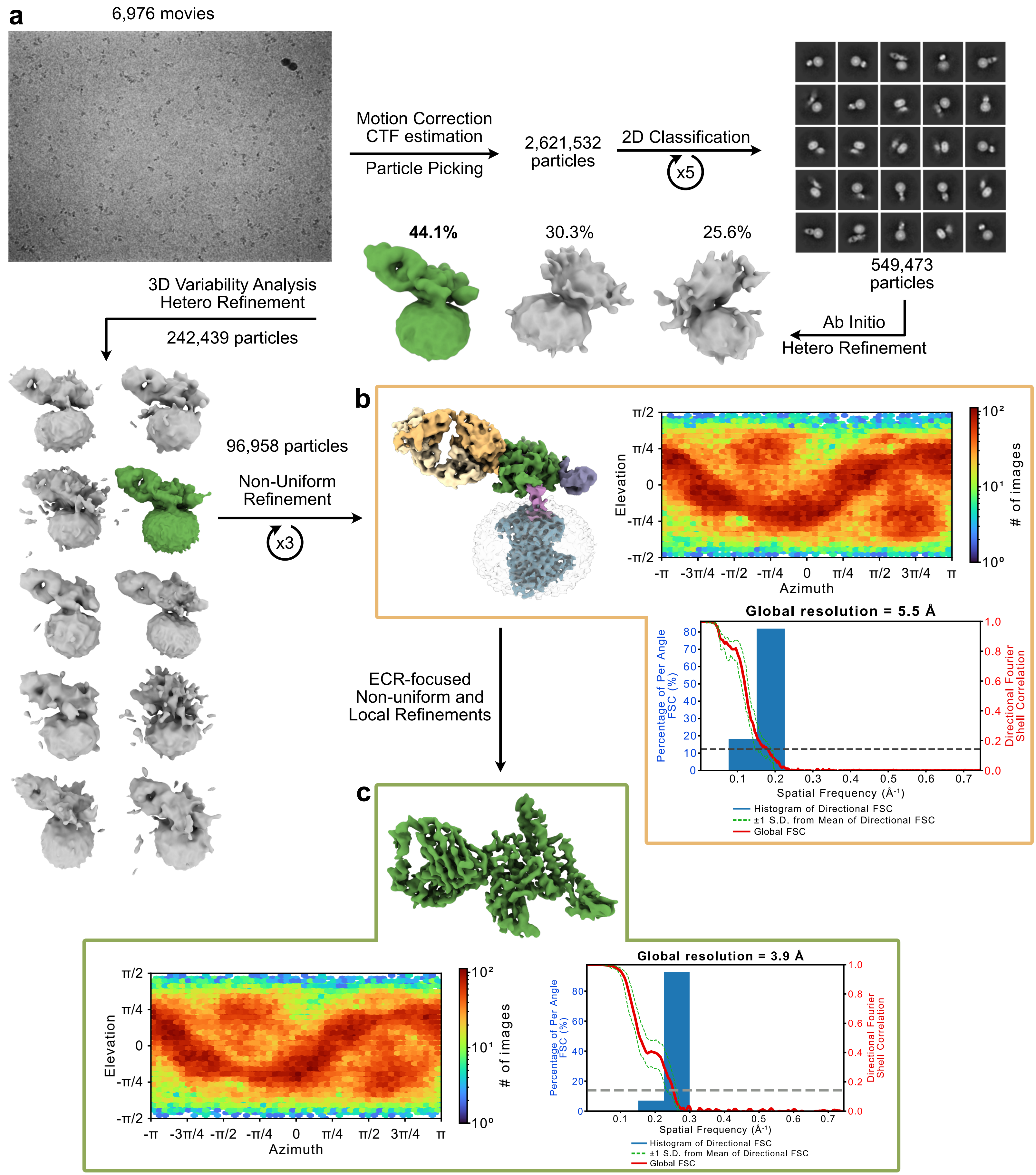


**Supplementary Fig. 2: Cryo-EM data processing of HormR+GAIN/7TM ADGRL3.**

**a** Flow chart of cryo-EM data processing and cryo-EM maps of ADGRL3 holoreceptor. **b** 5.5 Å map of ADGRL3 holoreceptor with a graph presenting angular distribution of particles used in the final 3D reconstruction and gold-standard Fourier shell correlation curves of the refinement. **c** Final 3.9 Å map of ADGRL3 HormR/GAIN domains in complex with sAB LK3. Below, a graph presenting angular distribution of particles used in the final 3D reconstruction and gold-standard Fourier shell correlation curves of the refinement.

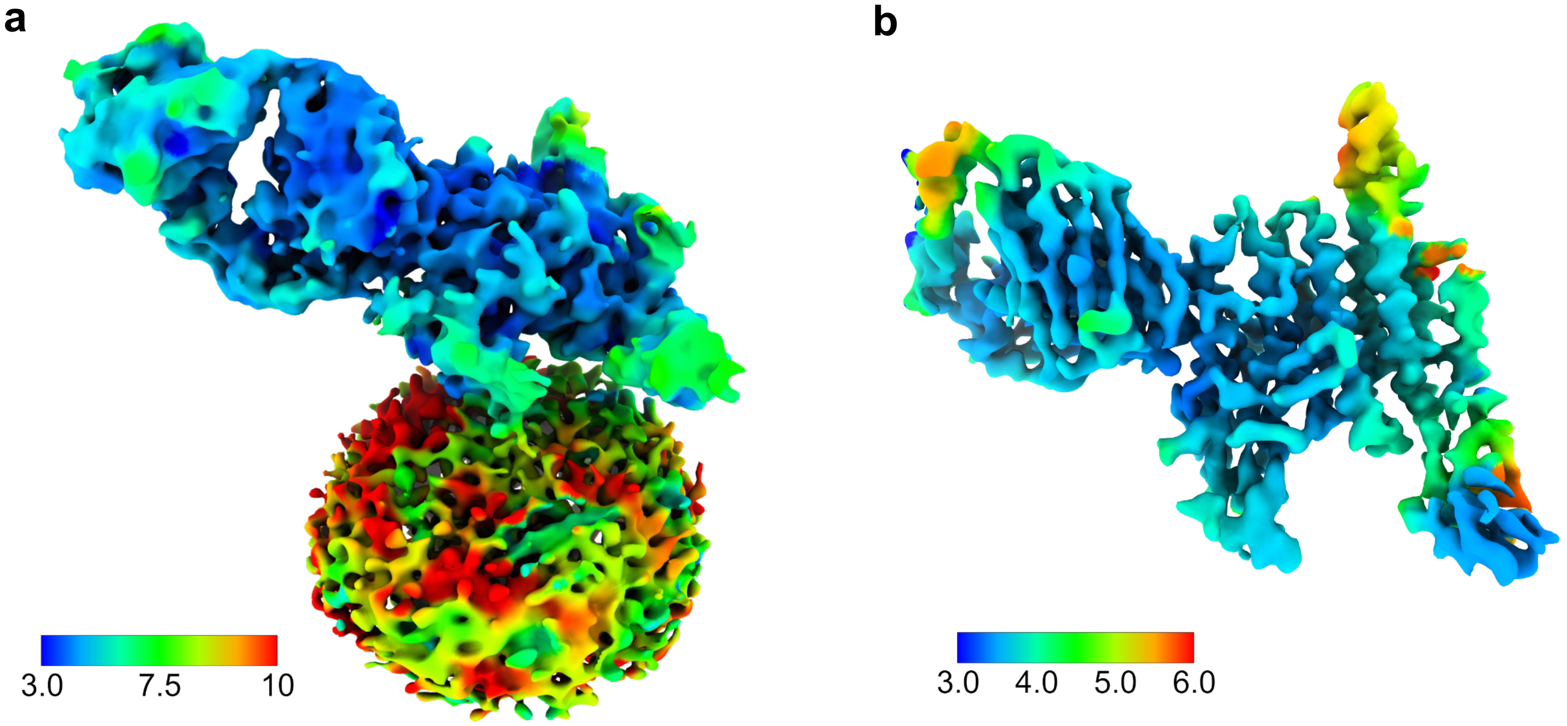


**Supplementary Fig. 3: 3D Cryo-EM maps of ADGRL3 colored by local resolution.**
A local resolution density maps of **a** ADGRL3 holoreceptor in detergent micelle and **b** HormR/GAIN domains in complex with sAB LK3, colored from blue (higher resolution) to red (low resolution).

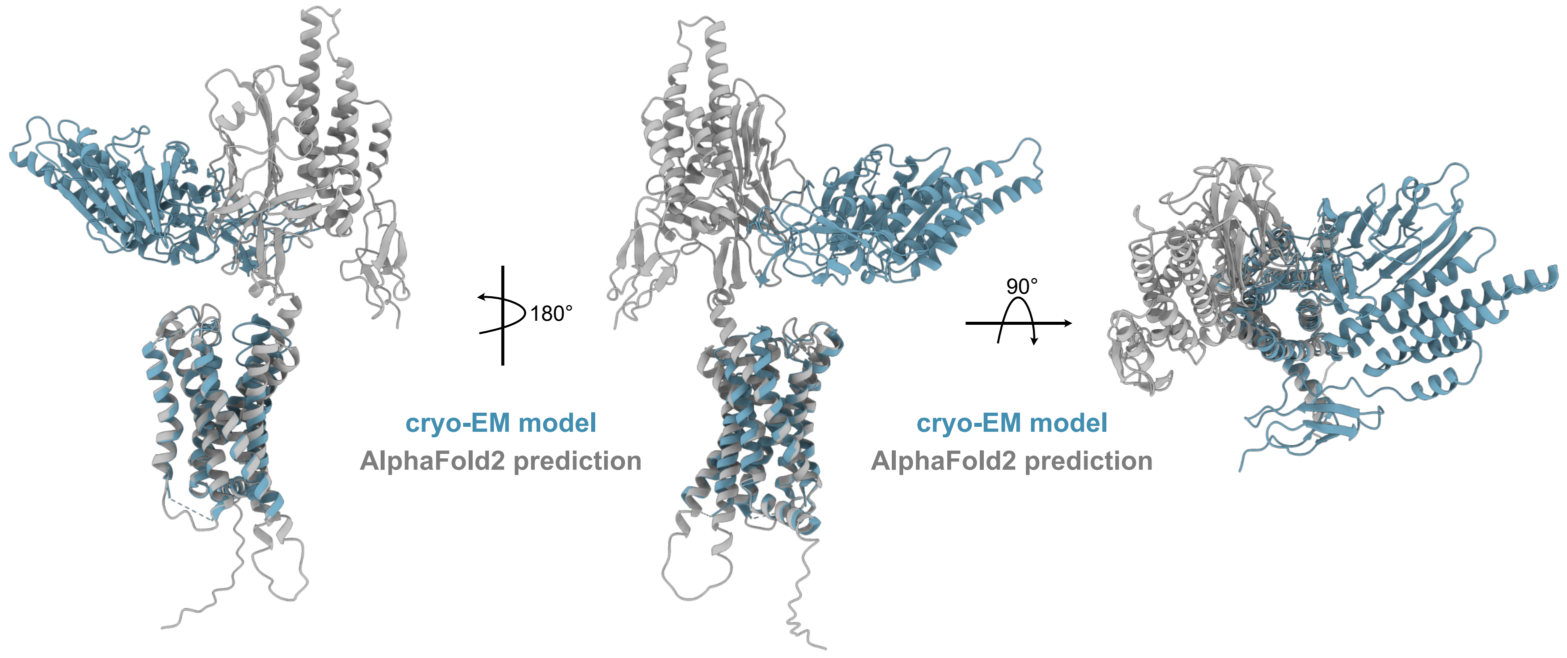


**Supplementary Fig 4: Comparison of ADGRL3 HormR+GAIN structure to AlphaFold2 prediction.**
Side and top views of ADGRL3 holoreceptor model based on low-resolution cryo-EM map (blue) with the AlphaFold2 prediction of the HormR+GAIN/7TM construct (gray) superimposed over 7TM domain.

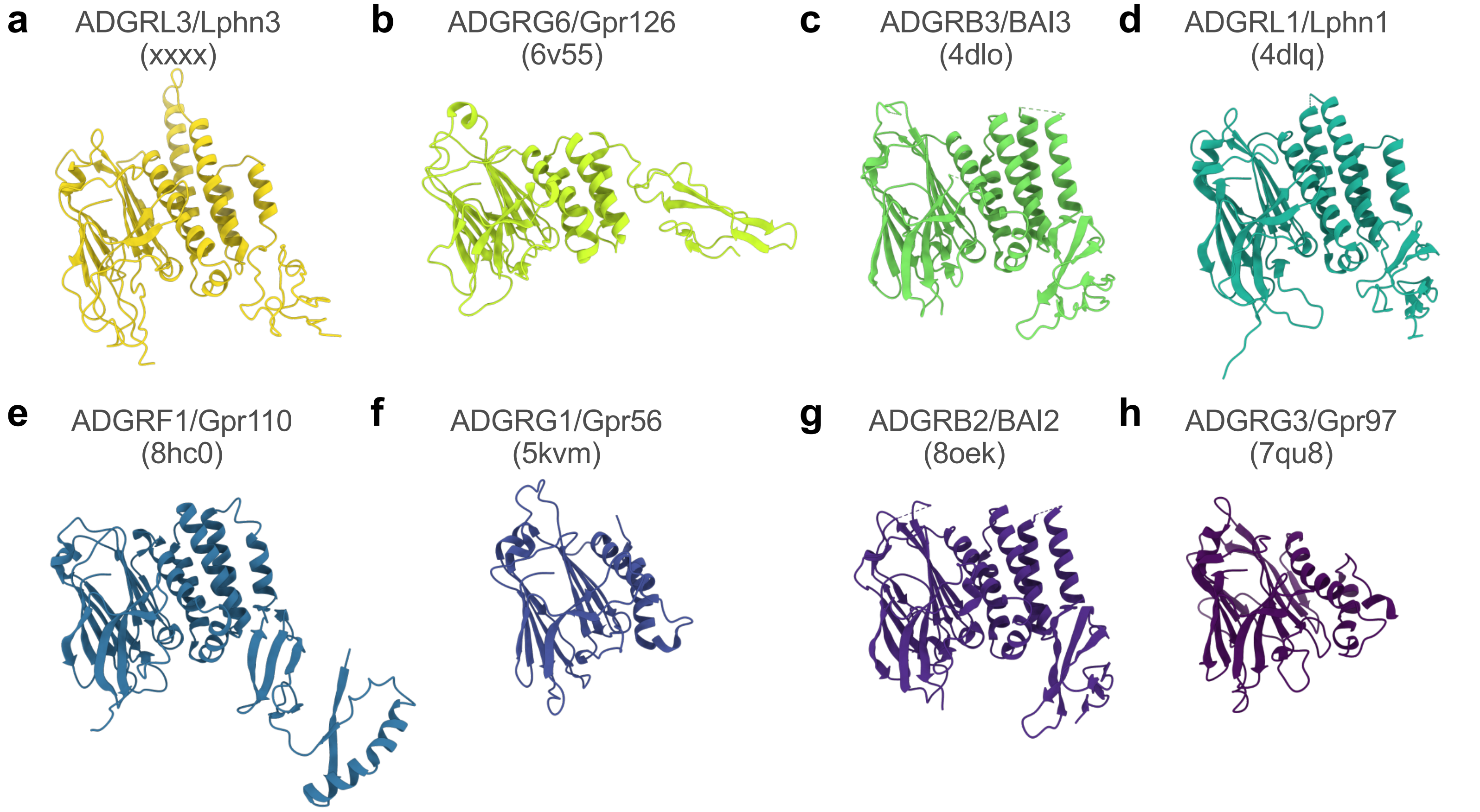


**Supplementary Fig 5: Comparison of ADGRL3 HormR+GAIN structure to available structures of GAIN domains of other adhesion GPCRs.**

GAIN domain structures of **a** ADGRL3/Lphn3 (PDB: xxxx), **b** ADGRG6/Gpr126 (PDB: 6v55), **c** ADGRB3/BAI3 (PDB: 4dlo), **d** ADGRL1/Lphn1 (PDB: 4dlq), **e** ADGRF1/Gpr110 (PDB: 8hc0), **f** ADGRG1/Gpr56 (PDB: 5kvm) **g** ADGRB2/BAI2 (PDB: 8oek) and **h** ADGRG3/Gpr97 (PDB: 7qu8), presented in the same orientation relative to ADGRL3, show conserved common fold of the domain.

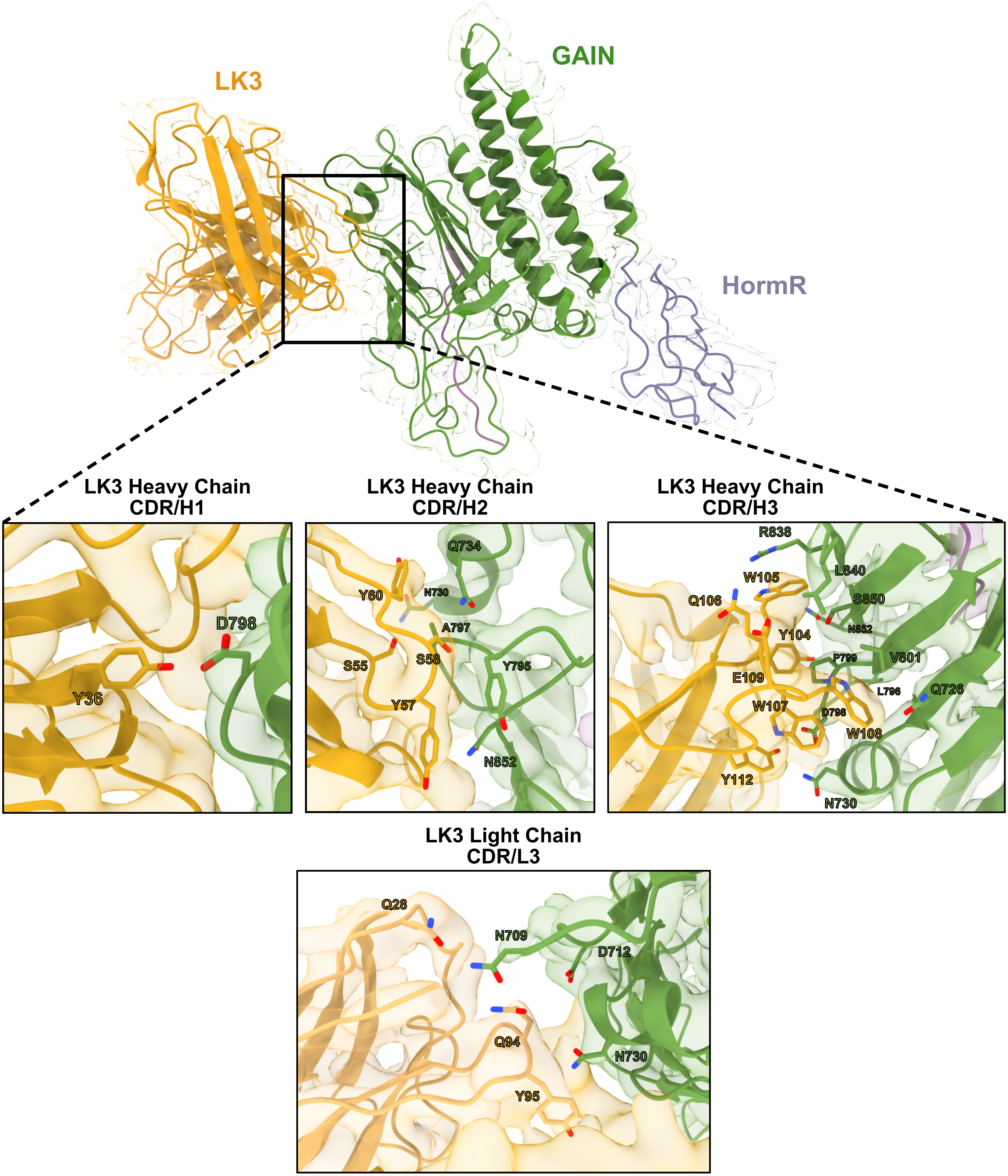


**Supplementary Fig 6: cryo-EM structure of LK3-bound HormR/GAIN domains of ADGRL3.**

Model of HormR/GAIN domains of ADGRL3 in complex with sAB LK3 fitted into the cryo-EM map; zoomed-in detailed views of crucial residues from LK3 complementarity-determining regions (CDRs, in yellow) making interactions with residues on GAIN domain of ADGRL3 (green) are shown as sticks.

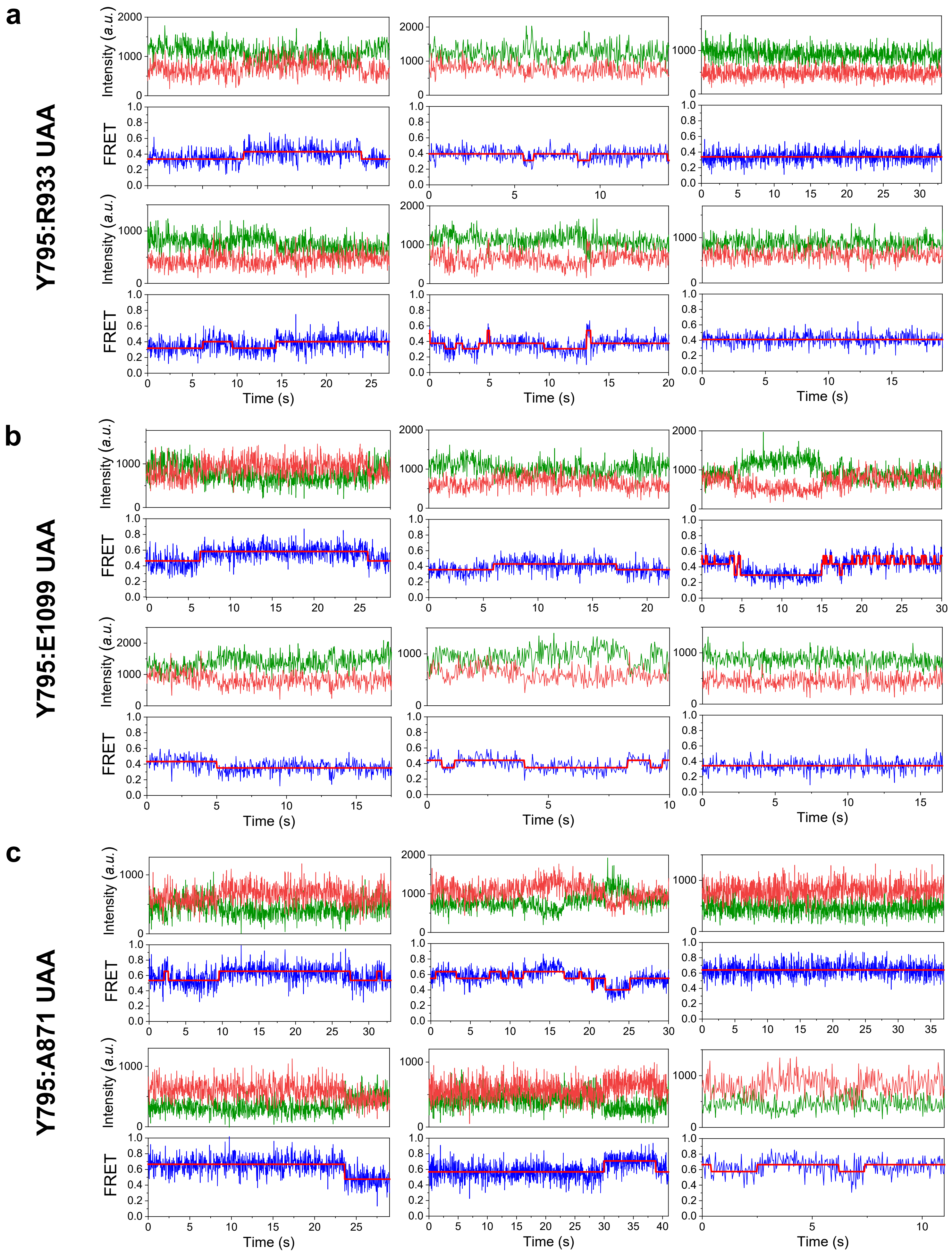


**Supplementary Fig. 7: Representative smFRET traces for different sensors.**

Representative smFRET traces of **a** Y795:R933 UAA **b** Y795:E1099 UAA **c** Y795:A871 UAA sensors showing traces with a single FRET state, two FRET states, and three FRET states before photobleaching

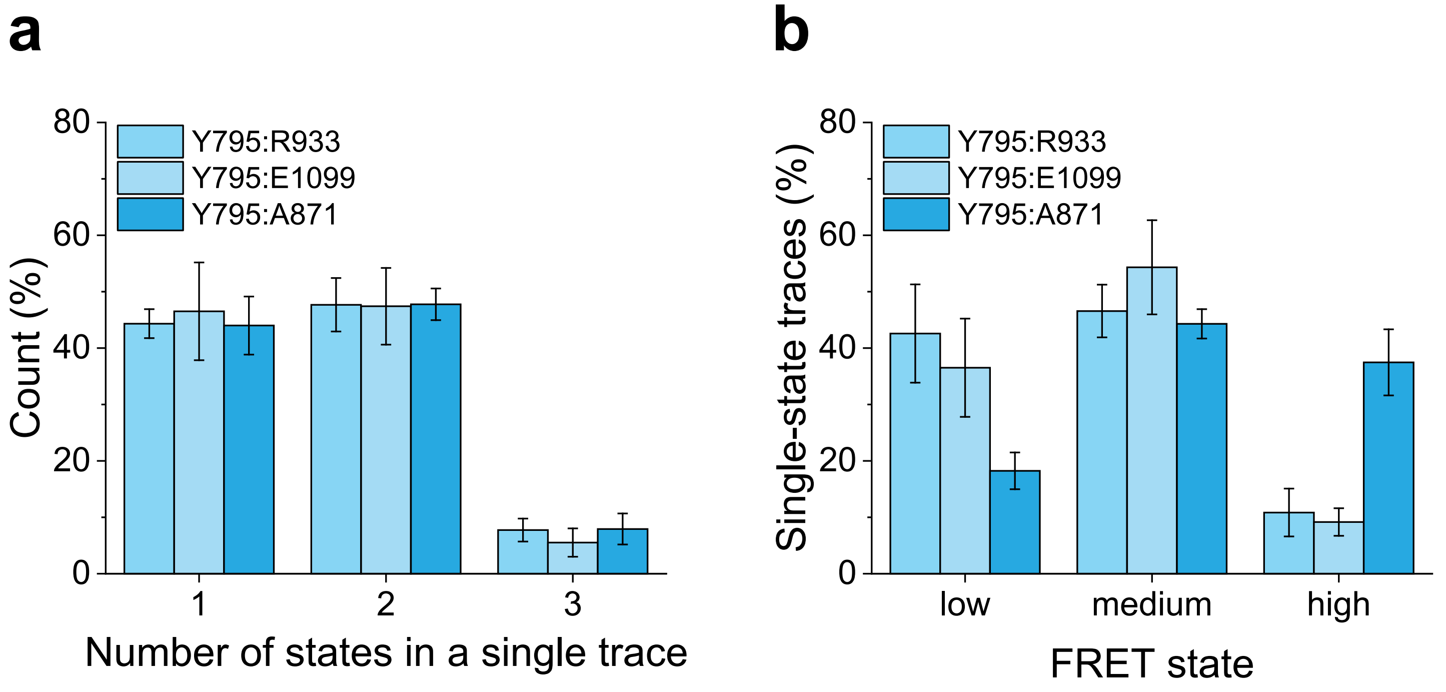


**Supplementary Fig. 8: smFRET states and occupancy quantification for sensors.**

**a** Quantification of the number of states in a single trace for each of the Y795:R933 UAA, Y795:E1099 UAA, Y795:A871 UAA sensors.

**b** Quantification of the occupancy of each FRET state for each of the Y795:R933 UAA, Y795:E1099 UAA, Y795:A871 UAA sensors.

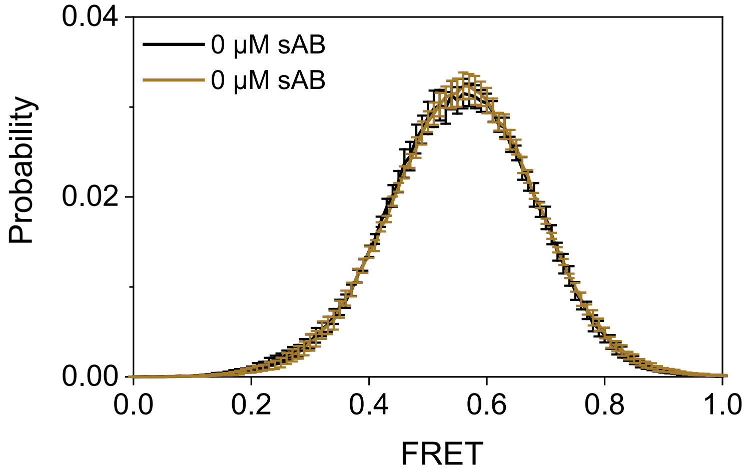


**Supplementary Fig. 9: smFRET population histogram in the absence and presence of 1 μM of non-specific synthetic antibody fragment.**

Control experiment with 1 μM of non-binding sAB showing no change in the smFRET histogram.

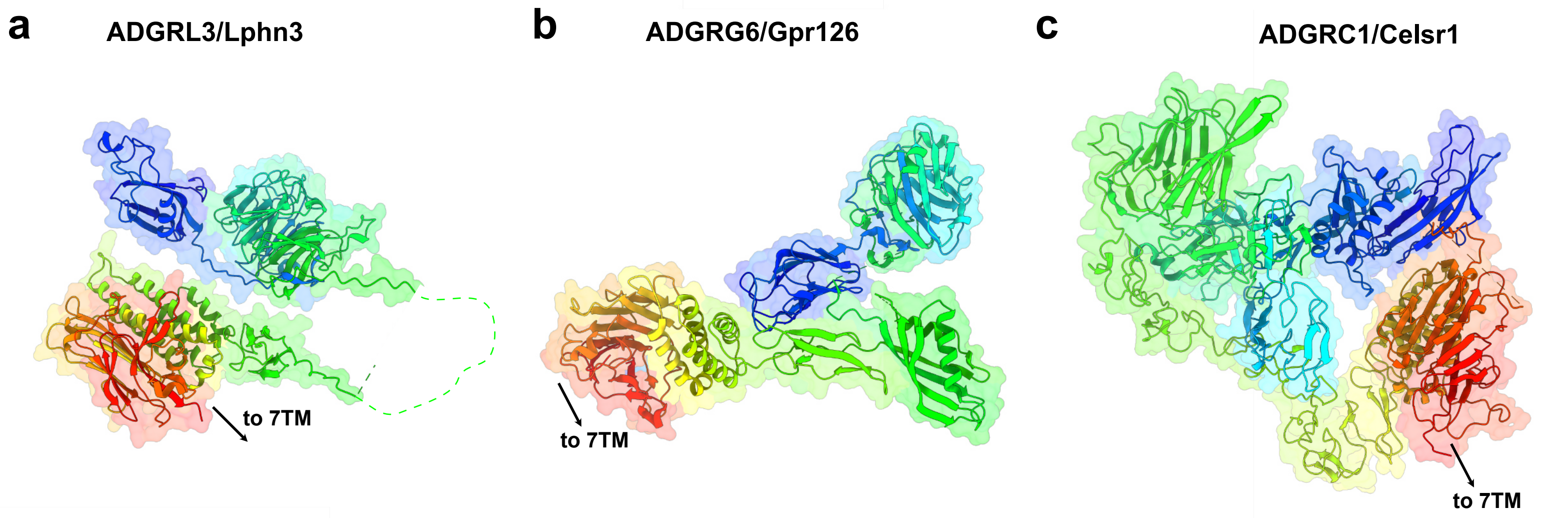


**Supplementary Fig. 10: aGPCRs adopt compact conformations of their extracellular regions.**

**a** AlphaFold prediction of ADGRL3 full-length ECR conformation. **b** X-ray crystallography structure of FL ECR of ADGRG6/Gpr126 (PDB: 6v55). **c** cryo-EM structure of C-terminal fragment of ECR of ADGRC1/CELSR1. Presented models, colored from blue (N-terminus) to red (C-terminus), show that ECRs of aGPCRs can form a compact multidomain modules.
